# Serine Racemase mediates Subventricular Zone Neurogenesis via Fatty acid Metabolism

**DOI:** 10.1101/2022.06.09.495488

**Authors:** Robin Roychaudhuri, Hasti Atashi, Solomon H. Snyder

## Abstract

The adult subventricular zone is one of the two neurogenic niches that continuously produce newborn neurons. Here we show that serine racemase (SR), an enzyme that catalyzes the racemization of L-serine to D-serine and vice versa, affects neurogenesis in the adult SVZ by controlling *de novo* fatty acid synthesis. Complete and conditional deletion of SR in nestin precursor cells lead to diminished neurogenesis in the SVZ. Nestin-cre+ mice showed reduced expression of fatty acid synthase and its substrate malonyl CoA, which are involved in *de novo* fatty acid synthesis. Global lipidomic analyses revealed significant alterations in lipid subclasses in nestin-cre+ mice. Decrease in fatty acid synthesis was mediated by phospho acetyl CoA carboxylase that was AMPK independent. Both L and D serine treatment rescued defects in SVZ neurogenesis, proliferation and levels of malonyl CoA *in vitro*. Our work shows that SR affects adult neurogenesis in the SVZ via lipid metabolism.

## Introduction

Adult neurogenesis is the production of new neurons in the adult brain (Kempermann, 2006; Ming and Song, 2011; Tong and Alvarez-Buylla, 2014). These new neurons are formed in the sub ventricular zone (SVZ) of the lateral ventricles and the sub granular zone of the hippocampus. Newborn neurons in the SVZ migrate along the rostral migratory stream (RMS) and eventually differentiate into olfactory bulb (OB) interneurons feeding the precursor cell population. In the adult SVZ this process is regulated by diverse genes and growth factors (Lim and Alvarez-Buylla, 2016). Recent work has shown that adult neurogenesis in both the neurogenic regions depends on metabolic control of fatty acid synthesis (Knobloch, 2017; Knobloch et al., 2013). Defects in fatty acid metabolism play a crucial role in the quiescent and proliferative behavior of neural progenitor cells (Clemot et al., 2020; Knobloch et al., 2017). Serine Racemase (SR) is a pyridoxal phosphate (PLP) dependent enzyme that synthesizes D-serine from L-serine and vice versa (Wolosker et al., 1999a; Wolosker et al., 1999b). D-serine is known to function as a neurotransmitter during brain development (Schell et al., 1997). SR deficient mice show NMDA receptor hypofunction and are considered a model for schizophrenia (Balu et al., 2013; Basu et al., 2009). However, little is known of D-serine’s functions outside the NMDA receptor. D-serine regulates proliferation and differentiation of neural stem cells (NSCs) from postnatal mouse forebrain (Huang et al., 2012). D-serine has been implicated in adult hippocampal neurogenesis, with exogenous treatment of D-serine increasing the density of neural stem cell, transit amplifying progenitors and increased survival of newborn neurons (Sultan et al., 2013). We investigated the metabolic role of SR in NSCs from the adult SVZ. We hypothesized that SR may influence metabolism in the SVZ due to L and D serine synthesis in astrocytes and neurons respectively (Wolosker et al., 2017) (Ehmsen et al., 2013).

We generated conditional knockout of SR in nestin precursor cells (nestin-cre+) to determine its role in metabolism (Bernal and Arranz, 2018). Nestin-cre+ mice show defects in lipid metabolism and adult SVZ neurogenesis.

Using lipidomic analysis and biochemical approaches we show that nestin-cre+ mice SVZ have significant alterations in the different classes of lipids, decreased expression of fatty acid synthase (FASN) and malonyl CoA, a precursor for *de novo* fatty acid synthesis. Our work also shows that SR mediates neurogenesis in the adult SVZ via acetyl CoA carboxylase (ACC) activation. SR may regulate this metabolic pathway by regulating the availability of astrocytic L-serine and or neuronal D-serine in NSCs. These findings may have therapeutic implications in the treatment of disorders of adult neurogenesis like schizophrenia and Alzheimer’s disease (Le Douce et al., 2020; Moreno-Jimenez et al., 2019; Schoenfeld and Cameron, 2015; Yun et al., 2016).

## Results

### Lack of SR leads to defects in adult SVZ neurogenesis and proliferation

SR is a PLP dependent enzyme, which synthesizes D-serine in neurons from L-serine transported from astrocytes (Wolosker et al., 2017; Wolosker et al., 1999b). We determined levels of neurogenesis in the SVZ of adult WT and SR^-/-^ mice (**Figure 1A, 1B, Suppl Figures 1A, 6A**). We monitored BrdU incorporation in age matched WT and SR^-/-^ mice for 14 days (Doetsch et al., 1999). Our data showed significantly reduced incorporation of BrdU in SR^-/-^ mice compared to WT (**Figure 1B, 1C**). Quantitative analysis showed four-fold reduction in BrdU positive cells in SR^-/-^ mice relative to WT (**Figure 1C**). GFAP positive staining (mature astrocytes) was decreased in the SVZ of SR^-/-^ mice compared to WT (**Figure 1B**).

**Figure 1.**
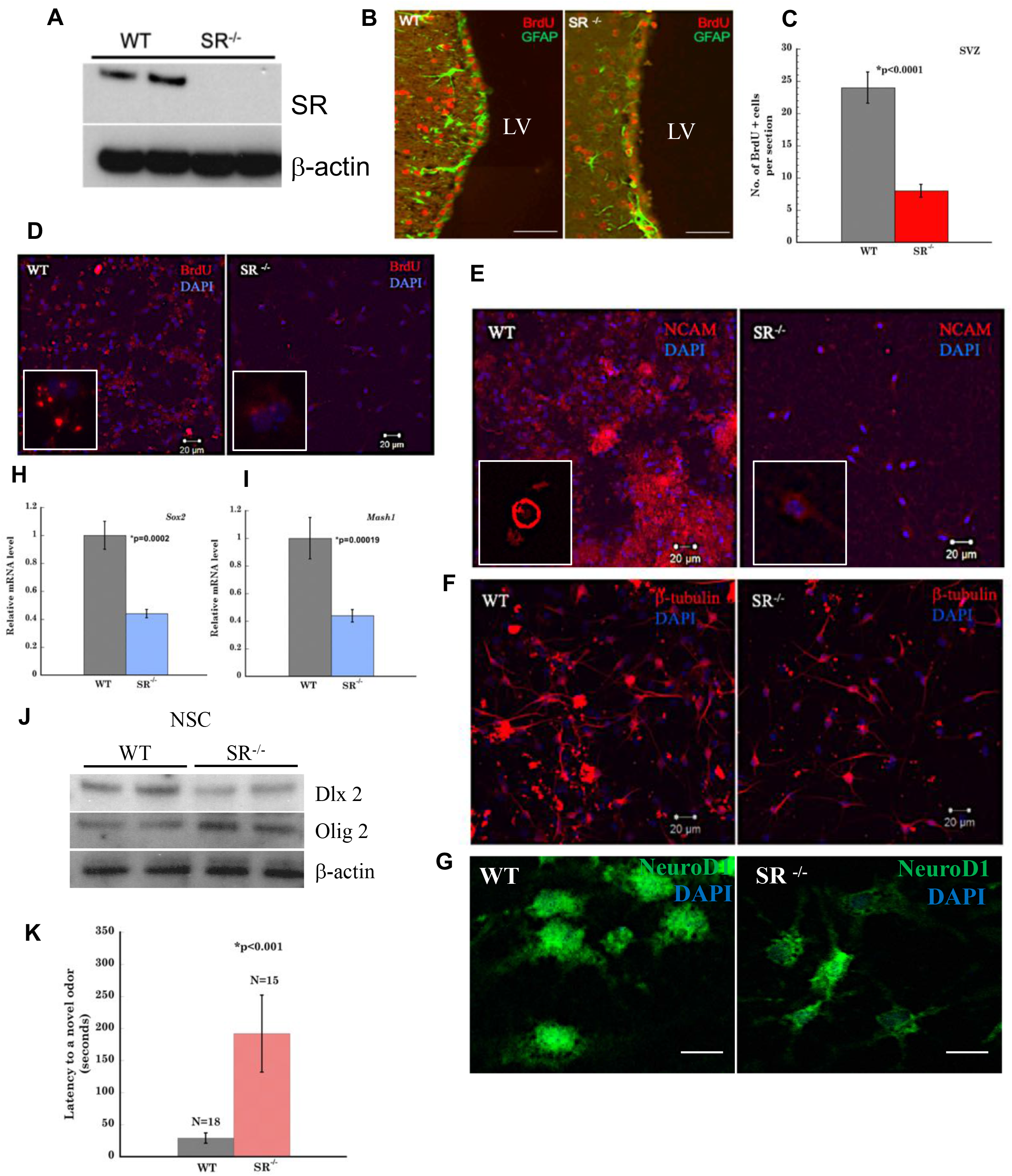
Lack of Serine Racemase leads to defects in adult SVZ neurogenesis and proliferation *in vivo*. (**A**) Expression of SR in WT and SR^-/-^ mice SVZ lysates. (**B**) Coronal section of WT and SR^-/-^ SVZ showing BrdU incorporation (red) and GFAP expression (green). Scale Bar= 200 µ. (**C**) Quantitative estimation of BrdU+ cells in the SVZ. (**D**) BrdU (10 µM for 18h) incorporation in NSCs isolated from WT and SR^-/-^ SVZ. Inset shows incorporation of BrdU in NSCs (red). Neuroblast assay showing expression of neuroblast markers (**E**) PSA-NCAM (inset shows PSA-NCAM expression in an isolated neuroblast), (**F**) β-III tubulin (**G**) NeuroD1 in NSCs isolated from SVZ of adult WT and SR^-/-^ mice 3-4 days after differentiation. mRNA levels of (**H**) *Sox2* (**I**) *Mash1* in olfactory bulb of WT and SR^-/-^ mice. Error Bars are SD. (**J**) Expression of Dlx2 and Olig 2 in NSC lysates from WT and SR^-/-^ mice. Actin was a loading control. (**K**) Olfactory behavior test in age matched WT and SR^-/-^ mice. N indicates the number of mice. p value indicates statistical significance (student’s t-test).

The neuroblast assay serves as an *in vitro* model of neurogenesis (Azari and Reynolds, 2016; Azari et al., 2012). An adherent monolayer culture of differentiating NSCs comprises a layer of astrocytes above which circular colonies of neuroblasts are present following induction of differentiation. To determine neurogenesis *in vitro*, we performed the neuroblast assay. NSCs were isolated from the SVZ of age matched adult WT and SR^-/-^ mice and grown as adherent monolayer cultures for 10-12 days (Walker and Kempermann, 2014). After 4 days incubation in media with reduced and eventually no growth factors, neuroblasts were observed above a layer of astrocytes (**Suppl Figure 2A**) (Azari and Reynolds, 2016). We examined BrdU incorporation in NSCs as an indicator of cell division. BrdU incorporation in NSCs from WT and SR^-/-^ SVZ showed four fold lower incorporation in SR^-/-^ compared to WT (**\***p=0.0084) (**Figure 1D and Suppl Figure 2E**).

Neuroblasts (type A cells) formed from NSCs migrate rostrally and differentiate to olfactory bulb interneurons. We determined expression of polysialylated neuronal cell adhesion molecule (PSA-NCAM), a marker for migrating SVZ neuroblasts. Neuroblasts from WT and SR^-/-^ SVZ were stained for PSA-NCAM, β-III tubulin (marker for immature neurons) and NeuroD1 (marker for neuroblasts). We observed significantly diminished expression of PSA-NCAM (**Figure 1E**) in SR^-/-^ NSC’s. Quantitative analysis shows approximately 5 fold fewer neuroblasts in SR^-/-^ mice compared to WT (**\***p<0.001) (**Suppl Figure 2B**). β-III tubulin staining in NSC’s was reduced in SR^-/-^ mice compared to WT (**Figure 1F**). Quantitative analysis showed 5 fold reduction of β-III tubulin positive staining in SR^-/-^ (**\***p<0.001) (**Suppl Figure 2C**). We observed a similar trend in expression of NeuroD1 (**\***p<0.001) (**Figure 1G and Suppl Figure. 2D**). We examined differentiation of the NSCs to astrocytes and neurons. GFAP expression was reduced in SR^-/-^ mice (**Suppl Figure 1C**). NeuN expression (marker for newborn post mitotic neurons) was also reduced in SR^-/-^ NSCs compared to WT (**Suppl Figure 1D**). Gene expression for markers of proliferation in the OB, *Sox2* and *Mash1* showed 2 fold reduction in SR^-/-^ mice (**Figures 1H and 1I**). To substantiate our findings, we determined expression of a proliferation marker Dlx2 in neural stem cells (WT and SR^-/-^). Our data show reduced expression of Dlx2 in isolated NSCs (**Figure 1J, Suppl Figure 8A**). We determined expression of Olig2, which is required for cell fate determination of motor neurons and oligodendrocytes (Wang et al., 2020). We observed 2 fold increased expression of Olig2 in SR^-/-^ NSCs (**Figure 1J, Suppl Figure 8B**) suggesting that deleting SR may favor oligodendrocyte differentiation. We determined expression of Ki67 in primary neurons derived from NSCs. SR^-/-^ neurons expressed 2 fold lower levels of Ki67 (**Suppl Figure 1B**). To determine the effects of SVZ neurogenesis on olfactory behavior, we performed olfactory behavior test in age matched WT and SR^-/-^ mice (N=15-18 mice per group) (Witt et al., 2009). In the novel odor test, we determined the amount of time the mice spent locaiting and sniffing the novel odor. The results showed significant difference (**\***p<0.001) with the WT mice taking shorter time to explore and sniff the novel odor compared to the SR^-/-^ mice that took approximately 8 times longer (**Figure 1K**). These data show that defects in SVZ neurogenesis in SR^-/-^ mice affect olfactory function. Collectively these results show that SR^-/-^ mice have defects in adult SVZ neurogenesis and proliferation in vitro and in vivo.

### Conditional deletion of serine racemase leads to defects in adult SVZ neurogenesis and proliferation

Nestin is a cytoskeletal protein expressed widely in the lateral wall of the lateral ventricle. It is a marker for neural progenitor cells (Bernal and Arranz, 2018; Wiese et al., 2004). We generated conditional knockout of SR in nestin precursor cells (nestin-cre+) to elucidate the function of SR in neural progenitor cell development. Our rationale was two-fold first, D-serine the product of SR may control neuronal development and generation of newborn neurons (Schell et al., 1997). Secondly, neuronal D serine synthesis dependent on astrocytic L-serine may influence proliferation and metabolic homeostasis. SR expression was reduced in the brains of nestin-cre+ mice compared to age matched SR ^flox/flox^ (**Figure 2A**). We ascertained levels of neurogenesis in the SVZ of adult age matched SR ^flox/flox^ and nestin-cre+ mice by monitoring BrdU incorporation (Doetsch et al., 1999). BrdU imaging in the SVZ showed 1.5 fold reduced incorporation in nestin-cre+ mice compared to control (**Figures 1B, 1C**). Expression of Ki67, a proliferative marker for early phase of adult neurogenesis showed approximately two fold expression in the SVZ of nestin-cre+ mice compared to SR ^flox/flox^ (**Figures 2D, 2E**) (Kee et al., 2002).

**Figure 2.**
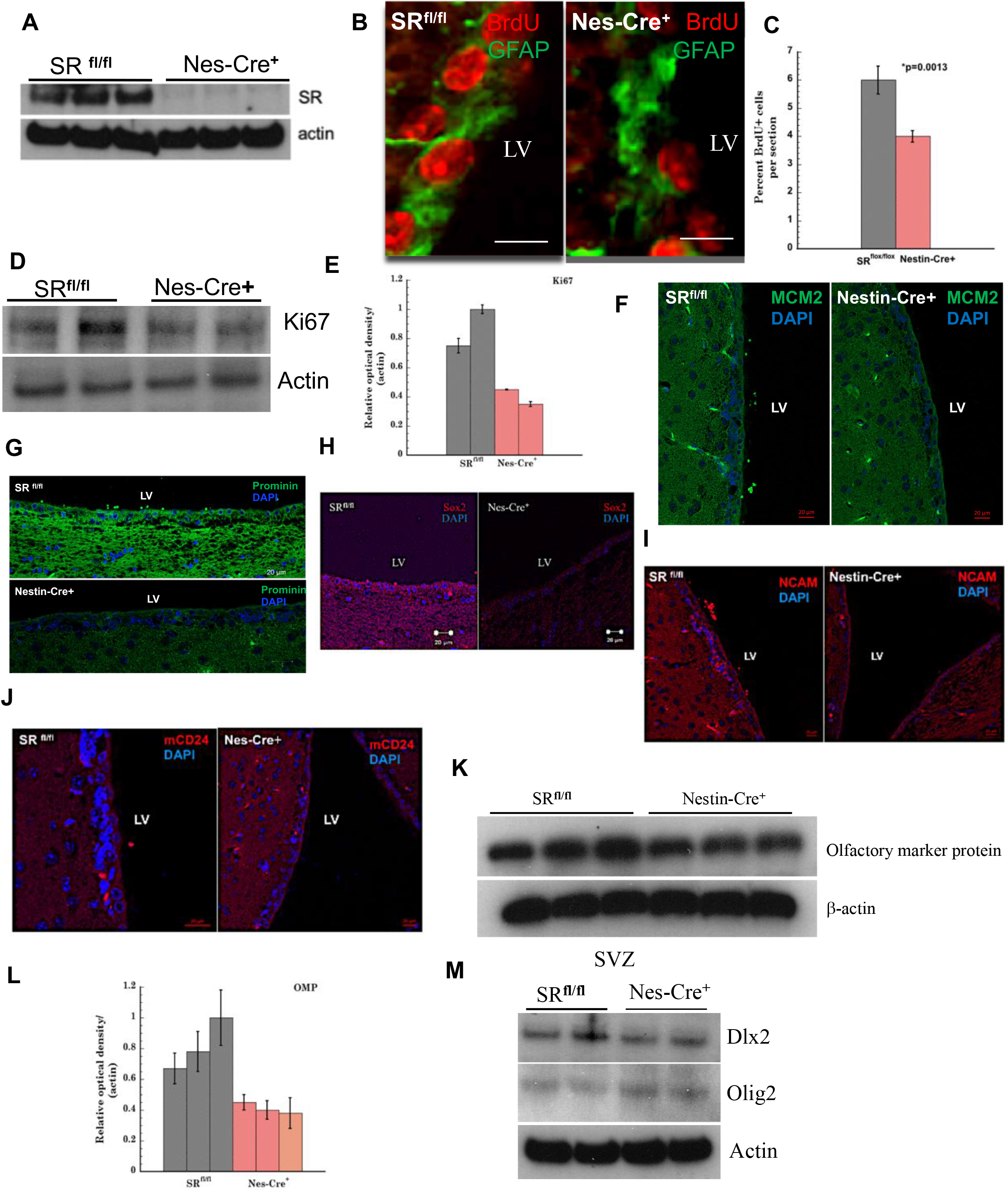
Conditional deletion of SR leads to defects in adult SVZ neurogenesis and proliferation. (**A**) Expression of SR in SR ^flox/flox^ and nestin-cre+ mice brain lysates. (**B**) BrdU incorporation (red) in the SVZ of SR ^flox/flox^ and nestin-cre+ mice and GFAP expression (green). Scale Bar=50 µ. LV (lateral ventricle). (**C**) Quantitative estimation of BrdU+ cells in the lateral ventricle of SVZ. (**D**) Western blot of SVZ lysates of SR ^flox/flox^ and nestin-cre+ mice showing expression of Ki67. (**E**) Densitometric analysis of blot in panel D. Expression of NSC and proliferation markers (**F**) MCM2 (**G**) Prominin (**H**) Sox2 (**I**) NCAM (**J**) mCD24 in the lateral ventricle of SVZ of adult SR ^flox/flox^ and nestin-cre+ mice. Scale Bar= 20 µ. (**K**) Expression of olfactory marker protein (OMP) in SVZ lysates of SR ^flox/flox^ and nestin-cre+ mice. Blots are representative of 3 independent experiments and pooled samples from N=5-7 mice per genotype. Samples were run in triplicate. (**L**) Plot shows densitometry of the OMP blot. (**M**) Blot shows expression of Dlx2 and Olig 2 in SVZ lysates of SR ^flox/flox^ and nestin-cre+ mice. Blot is representative of 3 independent experiments. Each sample was run in duplicate. Actin was a loading control.

We monitored expression of DNA replication licensing factor MCM2 (marker for proliferation of mitotic progenitor cells) in the SVZ. We observed approximately 3 fold reduction of MCM2 expression along the lateral wall of the SVZ in nestin-cre+ mice compared to SR ^flox/flox^ (**Figure 2F, Suppl Figure 9F**). Prominin 1 is a marker for NSCs in the ependymal and sub ependymal wall of the adult SVZ (Faigle and Song, 2013). Prominin 1 expression was reduced four fold in the SVZ of nestin-cre+ mice (**Figure 2G, Suppl Figure 9G**). We determined expression of mCD24 that is known to regulate the proliferation of committed neuronal precursors in the SVZ and is a negative regulator of cell proliferation (Belvindrah et al., 2002; Doetsch et al., 1999). Nestin-cre+ mice showed 3 fold higher expression of mCD24 in the SVZ compared to SR ^flox/flox^ (**Figure 2J, Suppl Figure 6B**). Sox2 is a transcription factor essential for self-renewal and differentiation of neural progenitor cells (Pevny and Nicolis, 2010). We determined Sox2 expression in the lateral wall of the SVZ and found 3 fold lower expression in nestin-cre+ mice (**Figure 2H, Suppl Figure 9I**). We determined expression of PSA-NCAM and observed 6 fold lower expression in the lateral wall of SVZ of nestin-cre+ mice (**Figure 2I, Suppl Figure 9H**). We also determined its expression along the RMS, and observed reduced expression in the nestin-cre+ mice (**Suppl Figure 5B**).

We determined expression of olfactory marker protein (OMP) present on olfactory receptor neurons originating from the SVZ. OMP expression was reduced 2 fold in nestin-cre+, mice suggesting that defects in adult SVZ neurogenesis affect olfactory function (**Figures 2K, 2L**). We observed reduced expression of proliferation marker Dlx2 in the SVZ of nestin-cre+ mice (**Figure 2M, Suppl Figure 8C**). We determined expression of Olig2 (cell fate determination of motor neurons and oligodendrocytes) (Wang et al., 2020). We observed increased expression of Olig2 in nestin-cre+ mice (**Figure 2M, Suppl Figure 8D**) suggesting that conditional deletion of SR may increase propensity for oligodendrocyte differentiation. Collectively our data show that nestin-cre+ mice have defects in SVZ neurogenesis and proliferation and support the data in SR^-/-^ mice **(Figure 1)**.

### SR alters L-serine and *de novo* fatty acid synthesis in the SVZ of Nestin-Cre+ mice

L-serine synthesized by astrocytes regulates neuronal development. It is a precursor for cysteine, sphingolipids, phosphatidylserine, phosphoglycerides, glycerides, nucleotides, D-serine and glycine (Furuya et al., 2000; Kalhan and Hanson, 2012; Mitoma et al., 1998; Reid et al., 2018). SR is the only enzyme known to predominantly synthesize D-serine (Wolosker et al., 2017; Wolosker et al., 1999b). We hypothesized that deletion of SR may have metabolic implications for neuronal development and maturation. L-serine synthesis is regulated by phosphoglycerate dehydrogenase (PGDH) and phosphoserine phosphatase (PSPH) via the phosphorylated pathway (Fell and Snell, 1988; Snell and Fell, 1990; Yamasaki et al., 2001). We determined expression of PGDH and PSPH in the SVZ of SR ^flox/flox^ and nestin-cre+ mice. Immunohistochemical data show decreased expression of both PGDH and PSPH along the lateral ventricle in nestin-cre+ mice (**Figure. 3A, Suppl Figures 9J, 9K**). Relative mRNA levels showed more than 50 % reduction in both enzymes in nestin-cre+ mice (**Figure 3B**). We determined expression at the protein level and observed decreased expression of both enzymes in the SVZ of nestin-cre+ mice (**Figure 3C**) supporting a consistent trend (Ehmsen et al., 2013). To determine if the presence of 0.4 mM L-serine in the culture media alters expression of PGDH and PSPH, we determined their expression in WT and SR^-/-^ NSCs. We found decreased expression of both PGDH and PSPH (**Figure 3E**) in the SVZ. FASN is a key enzyme that controls the synthesis of long chain fatty acids by synthesizing palmitate from acetyl and malonyl CoA. We determined expression of FASN in the SVZ that showed 3 fold reduction in FASN expression in nestin-cre+ SVZ (**Figure 3D, Suppl Figure 9E**). We also determined its expression in whole brain and SVZ and observed approximately 50 % lower FASN expression in nestin-cre+ mice (**Suppl Figures 2F, 2G**). To confirm our findings, we quantitatively estimated FASN levels in whole brain and SVZ lysates using ELISA. We found lower expression of FASN in nestin-cre+ brain and significantly lower levels in the SVZ (**Figures 3F and 3H**). These data strongly suggest that nestin-cre+ mice have defects in *de novo* fatty acid synthesis. Spot 14 (thyroid hormone responsive spot14) is a regulator of *de novo* lipogenesis and is involved in proliferation of adult NSCs (Knobloch et al., 2014). It negatively regulates malonyl CoA, an essential substrate for FASN in *de novo* lipogenesis. We determined expression of Spot14 in the SVZ of adult mice and observed greater expression of Spot 14 in nestin-cre+ mice (**Figure 3G**). These data suggest that increased Spot14 may play a role in FASN mediated regulation of SVZ neurogenesis in nestin-cre+ mice. Mig12 (<u>M</u>idline-1-Interacting <u>G</u>12 like protein) complexes with Spot14 to regulate acetyl CoA carboxylase (Knobloch et al., 2013; Park et al., 2013). We determined mRNA levels of *Mig12* in the SVZ and observed 10 fold higher expression in nestin-cre+ mice (**Figure 3I**). We estimated levels of malonyl CoA in the SVZ of age matched adult mice using quantitative ELISA. Our data show that levels of malonyl CoA in the SVZ of SR ^flox/flox^ were 1500 ng/ml, while in nestin-cre+ were 1200 ng/ml (*p=0.054 relative to SR ^flox/flox^ control) (**Figure 3J**). The reduced levels of malonyl CoA in nestin-cre+ mice suggest that *de novo* lipogenesis fueled by FASN and its substrate malonyl CoA is impaired.

**Figure 3.**
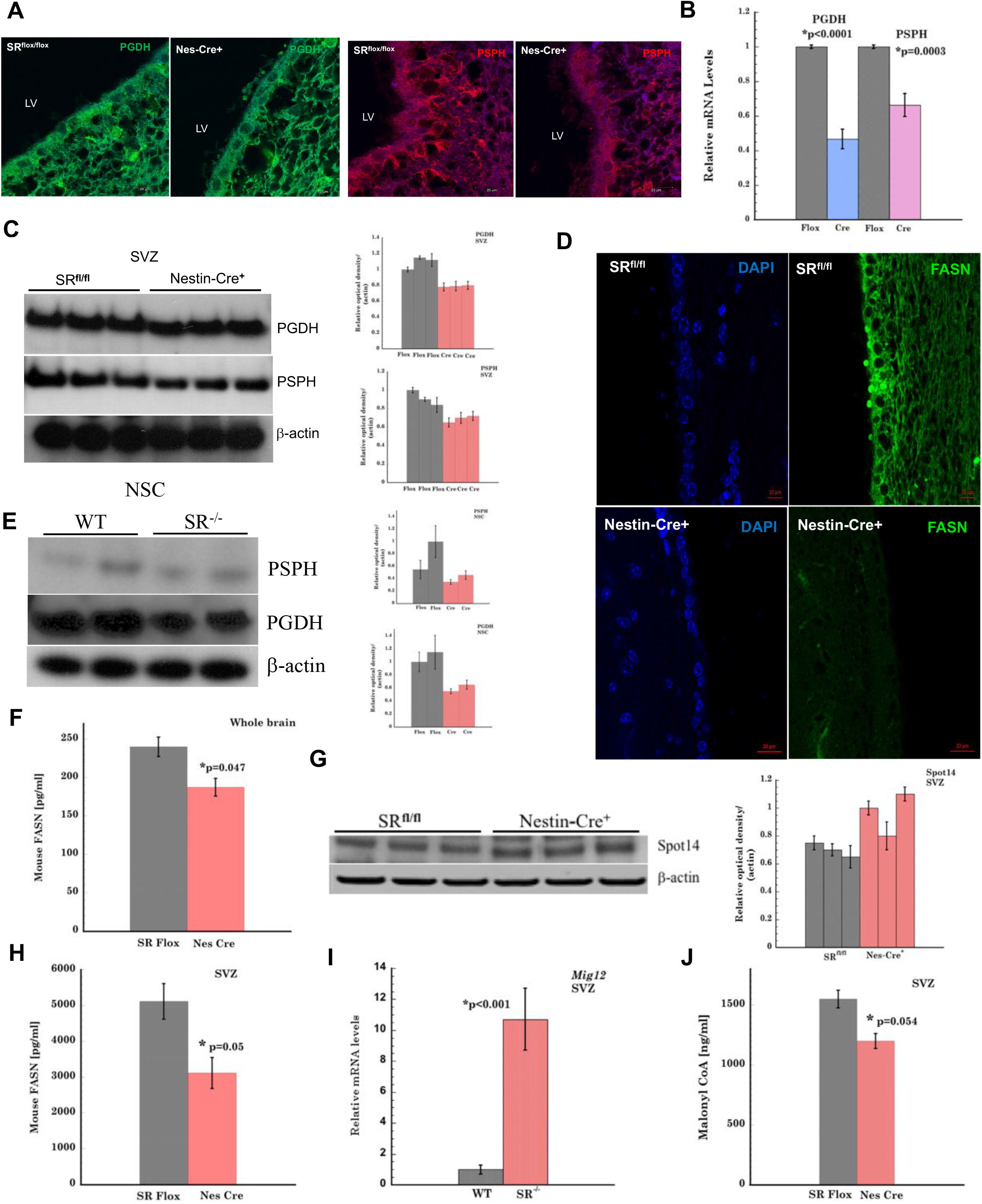
Serine Racemase alters L-serine and *de novo* fatty acid synthesis in the SVZ of nestin-cre+ mice. **(A)** Expression of L-serine synthesizing enzymes Phosphoglycerate dehydrogenase (PGDH) (green) and phosphoserine phosphatase PSPH (red) in the SVZ of SR ^flox/flox^ and nestin-cre+ mice. (**B**) Relative mRNA levels of PGDH and PSPH in the SVZ of SR ^flox/flox^ and nestin-cre+ mice. **(C**) Expression of PSPH and PGDH in SVZ lysates of SR ^flox/flox^ and nestin-cre+ mice. Panels on the right shows densitometric analysis of the blot on the left. (**D**) Expression of mouse fatty acid synthase (FASN) in the LV of the SVZ of adult, SR ^flox/flox^ and nestin-cre+ mice. DAPI is shown in the adjacent panel. Scale Bar=20 µ. (**E**) Expression of PSPH, PGDH in NSC lysates of WT and SR^-/-^ mice. Panels on the right shows densitometric analysis of the blots. (**F**) Quantitative estimation of mouse FASN expression in whole brain homogenates of SR ^flox/flox^ and nestin-cre+ mice by ELISA. Error bars refer to SD. **\*** indicates p=0.047 relative to SR ^flox/flox^ control. (**G**) Expression of Spot 14 in SVZ lysates of SR ^flox/flox^ and nestin-cre+ mice. Actin was a loading control. Panel on the right shows densitometric analysis of the blot. (**H**) Expression of mouse FASN in SVZ lysates of SR ^flox/flox^ and nestin-cre+ mice. Error bars refer to SD. **\*** indicates p=0.05 relative to SR ^flox/flox^ control. Data are representative of 3 independent experiments. (**I**) mRNA levels of Spot14 regulator Mig12 in SVZ of WT and SR^-/-^ mice. Data are representative of 3 independent experiments. (**J**) Estimation of malonyl CoA in the SVZ of SR ^flox/flox^ and nestin-cre+ mice by quantitative ELISA. **\*** indicates p=0.054 relative to SR ^flox/flox^ control. Data are representative of 3 independent experiments.

### Global lipidomic profile altered in the SVZ of adult SR ^flox/flox^ and Nestin-Cre+ mice

Recent evidence implicate lipids in regulating stem cell development and proliferation (Knobloch, 2017; Knobloch et al., 2013; Knobloch and Jessberger, 2017) (Lodhi et al., 2011). Our rationale for investigating the role of lipids is that serine metabolism regulates ceramide and sphingolipid metabolism as well as cell proliferation (Furuya et al., 2000; Gao et al., 2018; Kalhan and Hanson, 2012). We performed global lipidomic analysis in SVZ of age matched SR ^flox/flox^ and nestin-cre+ mice by lipid extraction followed by UPLC-MS/MS identification (**Figure 4A**). Volcano plot with fold change (mean nestin-cre+ / mean SR ^flox/flox^) ≥ 1.5 or ≤ 0.667 and p value adjusted for false discovery rate <0.05 as the criteria for significance, resulted in 184 lipids with fold change ≥ 1.5 (red) and 312 lipids with fold change ≤ 0.667 (blue) (**Figure 4B**). The different lipid classes were identified in 3 tiers of MS/MS analysis (**Figure 4C**). In tier 1(404 features with MS/MS match score threshold of 500), 2 (146 features with MS/MS match score threshold of 100) and 3 (remaining features in the lipidome database for putative mass match) with a mass error threshold of 5.0 mDa were identified. Out of 8648 unique features detected, 5262 features (60.6 %) were identified. Quality check (QC) was performed on all the samples by summation of intensities for all detected features in positive and negative ion mode. The summed intensities were consistent for extraction duplicates and injection triplicates for each sample as well as for the injection replicates for the QC sample showing good stability for data acquisition (**Suppl Figure 4B**). Internal standards were used to check for mass accuracy. Thirteen internal standards were detected in positive ionization and 12 internal standards in negative ionization (**Figure 4D**). The mass error was less than 2.8-3.0 ppm showing good mass accuracy. An average of 8215 ± 66 features per sample extraction were detected (**Suppl Figure 4C**). Heat map of the lipidomic analysis with the top 50 lipids (ranked by p value) is shown in **Figure 4E**. The heat map data show substantial downregulation of highly ranked lipids (by p value) in the SVZ of nestin-cre+ mice. Out of the top 50 lipids identified, 37 were downregulated and 13 upregulated in nestin-cre+ mice compared to SR ^flox/flox^. The identified lipids and their class are listed in **Suppl table 4** and in **Suppl Figure 11** respectively. Partial least squares-discriminant analysis with variable importance in prediction scores of 15 of the most important lipids (**Figure 4F, Table 1**) identified show a diverse group of lipids that are significantly altered in the SVZ. A caveat of nestin cre strain used in our study is the presence of metabolic phenotypes (Declercq et al., 2015; Harno et al., 2013). Nestin cre strain show lower levels of growth hormone that results in impaired insulin sensitivity and impaired glucose levels on a high fat diet. In order to rule out confounding effects, we estimated plasma glucose and insulin levels in age matched SR ^flox/flox^ and nestin-cre+ mice fed a normal diet. Our data show no difference in plasma glucose levels in SR^flox/flox^ and nestin-cre+ mice (**Figure 4G**). Total brain and SVZ insulin levels in SR ^flox/flox^ and nestin-cre+ mice also showed no differences (**Figures 4H, 4J**). We determined brain insulin levels in age matched WT and SR^-/-^ mice (background on which SR ^flox/flox^ mice were generated) to rule out any possible mice background effects. Our data showed no difference in brain insulin levels among WT and SR^-/-^ mice (**Figure 4I**). We determined expression of growth hormone in the SVZ of WT, SR^-/-^, SR ^flox/flox^ and nestin-cre+ mice. There was no change in growth hormone levels in WT and SR^-/-^ mice and a modest decrease in nestin cre mice compared to SR ^flox/flox^ **(Figure 4K)**. Our control experiments suggest that the metabolic differences in our work are due to conditional deletion of SR. The lipidomics data clearly highlight major differences among the different lipid classes in the neurogenic niche of the SVZ in nestin-cre+ mice.

**Figure 4.**
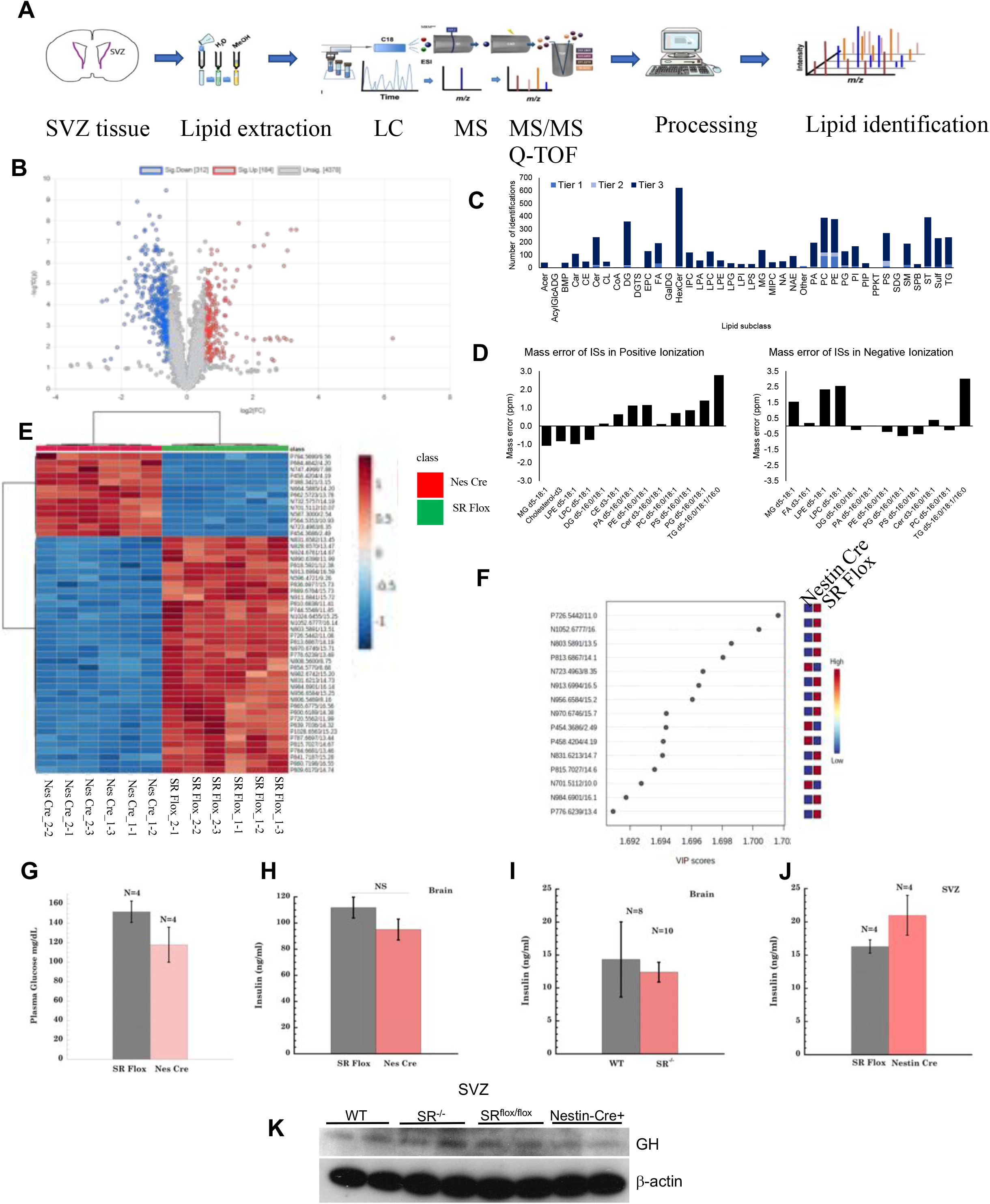
Altered global lipidomic profile in the SVZ of nestin-cre+ mice. (**A**) Schematic of lipid extraction followed by LC-MS/MS and subsequent lipid identification from the SVZ region of adult SR ^flox/flox^ and nestin-cre+ mice. (**B**) Parametric volcano plot showing fold change (calculated as mean (Nestin-Cre+) / mean (SR ^flox/flox^) against the p value adjusted for false discovery rate for each lipid. Blue circles indicate down regulation (312 lipids) while red indicates up regulation (184 lipids) of lipids. Unassigned lipids are shown in grey. (**C**) Lipid identification tier. In tier 1(404 features with MS/MS match score threshold of 500), 2 (146 features with MS/MS match score threshold of 100) and 3 (remaining features in the lipidome database for putative mass match) with a mass error threshold of 5.0 mDa were identified. (**D**) Mass error of internal standards in positive and negative ionization mode. Abbreviations for the different lipid categories are listed in Suppl Fig. 11. (**E**) Heat map of top 50 lipids ranked by p-value. Different lipids in the heatmap are identified in the Suppl table 4. (**F**) Plot shows the partial least squares-discriminant analysis (PLS-DA) variable importance in prediction (VIP) scores for 15 most important lipids. The identified lipids are listed in Suppl table 1. (**G**) Plasma glucose levels of SR ^flox/flox^ and nestin-cre+ mice estimated from tail bleed of mice randomly using glucose strips. (**H**) Insulin concentration (ng/ml) in whole brain homogenates of SR ^flox/flox^ and nestin-cre+ mice. Each genotype had an N=5-8 mice. (**I**) Insulin concentration (ng/ml) in whole brain homogenates of WT and SR^-/-^ mice. (**J**) Insulin concentration (ng/ml) in SVZ homogenates of SR ^flox/flox^ and nestin-cre+ mice Each genotype had an N=8-10 mice. Insulin levels were estimated using mouse insulin ELISA and low range insulin standard curve. Error Bars refer to SD. NS=Non Significant. (**K**) Blot of growth hormone in WT, SR^-/-^, SR ^flox/flox^ and nestin-cre+ SVZ lysates. Actin was a loading control. *<u>Note:</u>* SR Flox refers to SR ^flox/flox^ and Nes Cre refers to nestin-cre+ genotypes.

### Major Lipid classes altered in the SVZ of Nestin-Cre+ mice

Global lipidomic analysis in the SVZ of nestin-cre+ and SR ^flox/flox^ mice by UPLC-MS/MS showed significant alterations in the different classes of lipids. We plotted the data from the screen as fold change relative to SR ^flox/flox^ (significant). The different lipid classes were fatty acid conjugates, sphingomyelins, phosphatidylcholine, phosphatidic acid, ceramides, sterols, carnitine, tri, di and monoacylglycerols, hexosylceramides and lysophosphatidic acid. Fatty acid conjugates are polyunsaturated fatty acids with conjugate double bonds. Out of the 3 fatty acid conjugates identified, two showed modest decrease in nestin-cre+ compared to SR ^flox/flox^ while FA 12:0; O4 showed two fold higher levels in nestin-cre+ mice (**Suppl Figure 3A, Table 2**). L-Serine availability affects the synthesis of ceramides, sphingomyelin (sphinganine precursor), and phosphatidylserine (Esaki et al., 2015) (Gao et al., 2018). Our data show consistent reduction by > 50 % in all the sphingomyelin species identified (**Suppl Figure 3B, Table 2**). Most notable decrease was seen in sphingomyelin 38:1; O2 in nestin-cre+. Phosphatidylcholine is an abundant phospholipid in mammalian cell membranes. Phosphatidylcholine synthesis was significantly reduced (> 40 %) in the SVZ of nestin-cre+ mice (**Suppl Figures 3C, 4A and Table 2**). Phosphatidic acid (low abundance phospholipid) showed increases (≥2 fold) in PA 8:0_25:0, PA 36:1 and PA 19:2_19:2 and decrease (≥ 2 fold) in the remaining molecules in nestin-cre+ mice (**Suppl Figure 3D, Table 2**). Ceramide is a simple sphingolipid derived from sphingosine. All identified ceramides in our screen showed approximately 40 % or higher reduction in nestin-cre+ mice relative to SR ^flox/flox^ (**Suppl Figure 3E, Table 2**). Sterols are components of lipid rafts in membranes and play roles in membrane fluidity and signaling. Sterol expression was consistently increased above 3 fold in all the identified species with the exception of ST 22:1; O4 whose expression was down 10 fold in nestin-cre+ mice (**Suppl Figure 3F, Table 2**). Carnitine (a branched non-essential amino acid) expression was reduced by 50 % for 3 of the identified 4 species with the exception of CAR 24:1; O2 which was increased 2 fold in nestin-cre+ mice (**Suppl Figure 3G, Table 2**). Both tri and diacylglycerols (synthesized from fatty acyl CoA and glycerol 3 phosphate) were reduced by 50 % or more in nestin-cre+ mice with the exception of TG 70:3, TG 75:4 (**Suppl Figure 3H, Table 2**) and DG 34:1 and DG O-24:1 (**Suppl Figure 3I, Table 2**) that were increased (≥ 2.5 fold). Monoacylglycerols were increased 2 fold in nestin-cre+ mice (**Suppl Figure 3J, Table 2**). Hexosylceramides are sphingolipids with hexose ring attached to a ceramide, which is a major hub of the sphingolipid. Nestin-cre+ mice showed consistent downregulation (≥ 40%) in all the hexosylceramide molecules identified (**Suppl Figure 3K, Table 2**). Lysophosphatidic acid (a bioactive phospholipid produced during cell membrane synthesis) showed trends in both directions with LPA 30:2 and LPA 32:3 being reduced 2 fold and molecules LPA O-18:3, LPA O-22:3 and LPA O-26:3 increased more than 5 fold in nestin-cre+ mice (**Suppl Figure 3L, Table 2**). Our lipidomic screen shows clear trends towards reduction in most of the lipid classes in the SVZ of nestin-cre+ mice.

### Non AMPK dependent acetyl CoA carboxylase (ACC) control of fatty acid metabolism in Nestin-Cre+ SVZ

ACC is a regulator of *de novo* fatty acid synthesis (Garcia and Shaw, 2017; Steinberg et al., 2006). We observed increased activation of phosho-ACC (Ser79) in the SVZ of nestin-cre+ mice (**Figures 5B, 5C**). AMP-activated protein kinase (AMPK) is a cellular sensor of nutrient availability and ATP. It controls overall cellular lipid metabolism through direct phosphorylation of ACC1 and ACC2 at Ser 79, suppressing fatty acid synthesis and simultaneously promoting fatty acid oxidation by relieving suppression of carnitine palmitoyl transferase 1 (CPT1) (Gonzalez et al., 2020) (Garcia and Shaw, 2017). We observed more than 2 fold increased expression of AMPK (α) in SVZ of nestin-cre+ mice (**Figures 5A, 5E**). Increased activation of AMPK (α) is known to phosphorylate ACC and inhibit the synthesis of malonyl CoA (Garcia and Shaw, 2017; Steinberg et al., 2006). Expression of p-AMPK (Thr172) levels, were surprisingly 2 fold lower in nestin-cre+ SVZ compared to SR ^flox/flox^, suggesting a non-AMPK dependent phosphorylation of ACC (**Figures 5B, 5D)**. Based on the observed trends in *de novo* fatty acid synthesis, we determined expression of key genes in the synthesis and β-oxidation. Citrate lyase (CL) converts citrate from carbohydrate metabolism to acetyl CoA. We observed an 8 fold increase in *CL* mRNA expression in nestin-cre+ mice (**Figure 5G**). Cpt1a is a key rate-limiting enzyme in long chain fatty acid oxidation and its activity is negatively regulated by malonyl CoA. We observed approximately 500 fold increase in *Cpt1a* mRNA expression in nestin-cre+ SVZ (**Figure 5G**). Acc1 catalyzes the biotin dependent irreversible carboxylation of acetyl CoA to malonyl CoA, which serves as a key precursor in fatty acid synthesis. Our data showed 8 fold downregulation of *Acc1* in the SVZ of nestin-cre+ mice (**Figure 5G**). We next determined mRNA levels of key genes involved in β-oxidation. Malonyl CoA decarboxylase (MCD) that converts malonyl CoA to acetyl CoA was downregulated by 50 % in nestin-cre+ mice (**Figure 5F**). Hydroxyacyl CoA (HA CoA) dehydrogenase is involved in β-oxidation of hydroxyacyl CoA to oxoacyl CoA. In the SVZ of nestin-cre+ mice, the expression of *HA CoA* was elevated 30 fold compared to SR ^flox/flox^ (**Figure 5F**). Enoyl CoA hydratase (ECH) catalyzes the hydration of double bond between the second and third carbon atoms on 2-trans/cis enoyl-CoA thioester to produce β-hydroxyacyl CoA thioester. We observed > 8 fold expression of *ECH* in the SVZ of nestin-cre+ mice (**Figure 5F**). Collectively, these data suggest that SR may alter serine synthesis and availability compromising the *de novo* fatty acid synthesis pathway due to lower levels of malonyl CoA, FASN and downregulation of enzymes in fatty acid synthesis via activation of p-ACC by an unknown kinase. A concomitant increase in β-oxidation may compensate for the production of energy in the SVZ of nestin-cre+ mice (**Figure 5H**).

**Figure 5.**
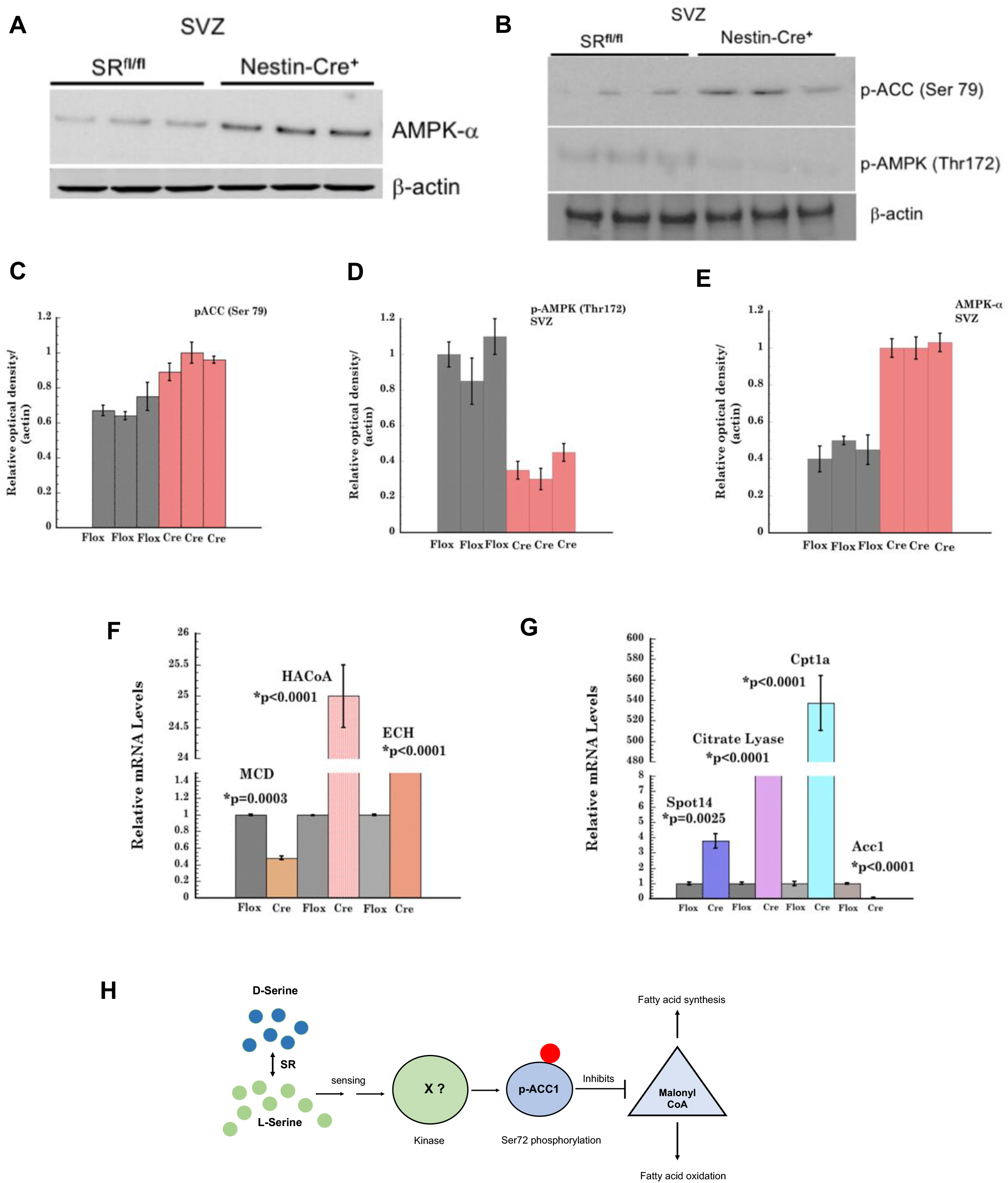
Expression of AMPK-α, phospho-AMPK, phospho-ACC and fatty acid metabolism enzymes. (**A**) Expression of AMPK-α in the SVZ of SR ^flox/flox^ and nestin-cre+ mice. (**B**) Expression of p-ACC (Ser79) and phospho-AMPK (Thr172) in the SVZ of SR ^flox/flox^ and nestin-cre+ mice. Actin was a loading control. (**C**) Densitometric analysis of p-ACC (Ser79) blot. (**D**) Densitometric plot of p-AMPK-α blot. (**E**) Densitometric plot of AMPK-α blot. (**F**) Relative mRNA levels of fatty acid oxidation genes Malonyl CoA decarboxylase (*MCD*), Hydroxy acyl CoA (*HACoA*) and Enoyl CoA hydratase (*ECH*) in the SVZ of mice. (**G**) Relative mRNA levels of *Cpt1a* (carnitine palmitoyl transferase 1a), *Acc1* (acetyl coA carboxylase 1), *Spot 14* and *Citrate lyase* in the SVZ of mice. Data are mean of 3 independent experiments from 3 different preparations of RNA. Error Bars are SD.(**H**) Schematic of SR mediated phosphorylation of ACC by a kinase. *<u>Note:</u>* Flox refers to SR ^flox/flox^ and Cre refers to nestin-cre+ genotypes.

### L and D Serine rescue defects in adult SVZ neurogenesis and proliferation in vitro

L-serine is an essential amino acid involved in sphingolipid synthesis and possibly in AMPK activation leading to phosphorylation of ACC and inhibition of malonyl CoA in *de novo* fatty acid synthesis (Garcia and Shaw, 2017). In order to determine if defects in adult SVZ neurogenesis and proliferation in nestin-cre+ and SR^-/-^ mice could be rescued by treatment with L-serine and D-serine *in vitro*, we performed rescue experiments in adherent NSC cultures treated with exogenous L-serine (0.6 mM) and D-serine (0.1 mM) separately and in an equimolar racemic mixture (0.1 mM each). Since neurobasal media contains basal L-serine concentration of 0.4 mM, and no D-serine, we used L-serine at a concentration of 0.6 mM to determine its effect. Control conditions comprised media containing 0.4 mM L-serine but without the exogenous L and or D serine. We determined incorporation of BrdU in WT and SR^-/-^ NSCs. SR^-/-^ NSCs show significantly reduced incorporation of BrdU compared to WT (**Figure 6A**). However, addition of exogenous L-serine and D-serine increased the number of proliferating cells incorporating BrdU in SR^-/-^ NSCs (**Figures 6B, 6C, 6F**). L-serine treatment showed higher levels of BrdU incorporation in SR^-/-^ NSCs compared to D-serine. We also determined rescue of Sox2 in NSCs (**Figure 6D**). We observed lower expression of Sox2 in SR^-/-^ NSCs compared to WT in control condition. However, upon treatment with equimolar mixture of L and D serine, we observed increased expression of Sox2 in both WT and SR^-/-^ NSC (**Figure 6D, Suppl Figure 9B**). NeuN is a marker for newborn neurons. We observed decreased NeuN expression in SR^-/-^ control NSCs, but upon treatment with equimolar mixture of L and D serine, NeuN expression was higher than WT control. We also observed that treated WT NSCs showed increased NeuN expression compared to control, indicating that both L and D serine play a role in the formation and maturation of newborn neurons (**Figure 6E, Suppl Figure 9A**). Nestin-cre+ mice showed increased expression of AMPK-α (**Figure 5B**). To determine if replenishing L and or D serine has an effect on expression of AMPK-α, we determined its expression in a rescue experiment. SR^-/-^ NSCs growing in presence of L and D serine (0.6 mM and 0.1 mM respectively) showed decreased expression of AMPK-α compared to untreated controls, indicating that AMPK-α expression is regulated by L and or D-serine. Interestingly, D-serine treatment appeared to have a more potent effect on AMPK-α expression compared to L-serine (**Figure 6G**).

**Figure 6.**
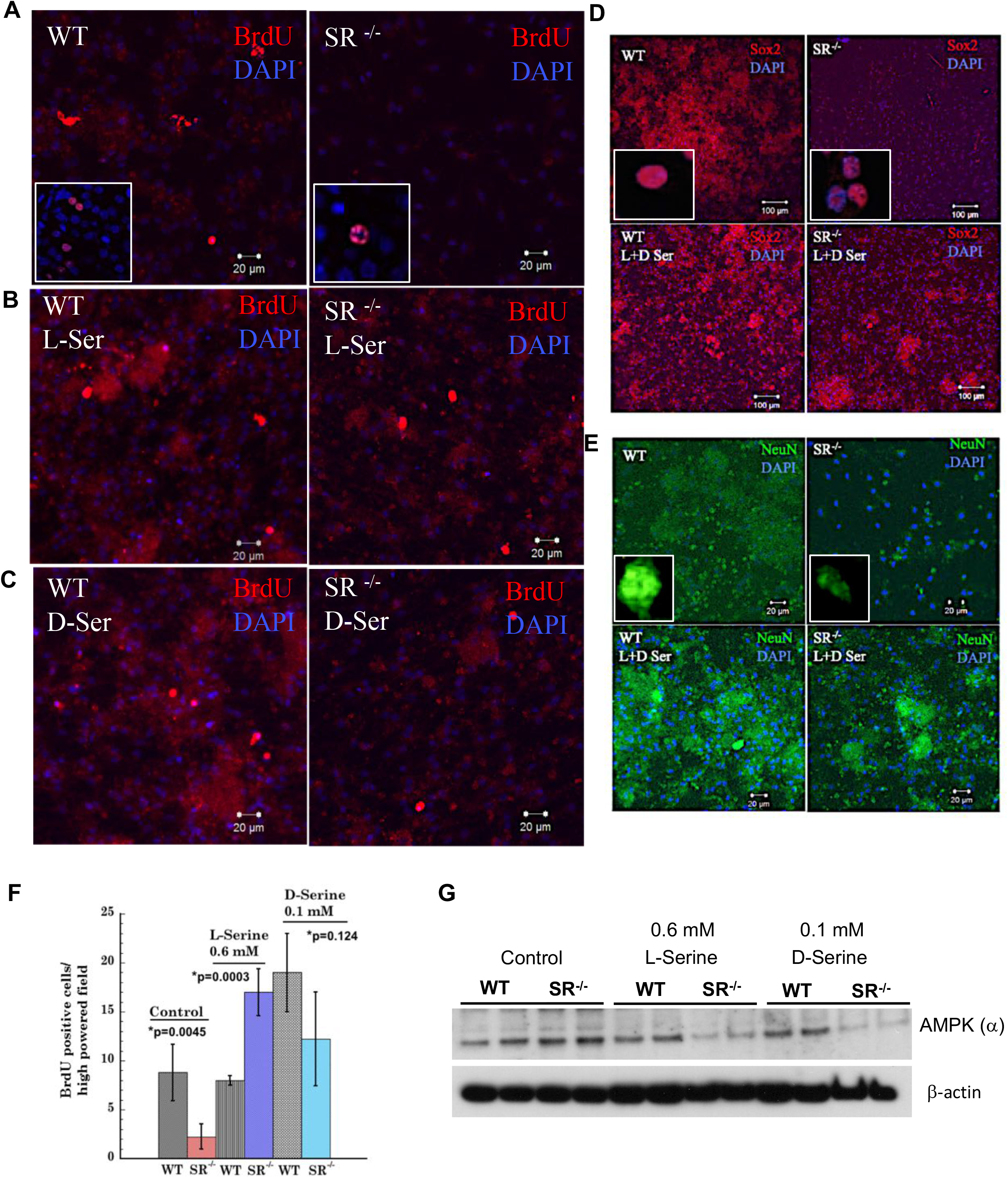
L and D serine rescue defects in SVZ neurogenesis and proliferation in vitro. WT and SR^-/-^ NSCs were grown in 0.6 mM L-serine and 0.1 mM D-serine in media. After induction of differentiation, the cells were fixed, permeabilized and stained. **(A)** BrdU incorporation in control WT and SR^-/-^ NSCs treated with 10 µM BrdU for 18 h at 37°C. **(B)** BrdU incorporation in cells treated with 0.6 mM L-serine **(C)** BrdU incorporation in cells treated with 0.1 mM D-serine. **(D)** Expression of Sox2 in control (top) and NSCs treated with equimolar mixture of L and D serine (100 µM each) (lower panel). **(E)** Expression of NeuN in differentiating neurons in control (top) and in presence of a mixture of L and D serine (bottom). DAPI is indicated in blue. Scale Bar=20 µ. **(F)** Number of BrdU positive cells counted per high powered field in NSC’s from WT and SR^-/-^ mice SVZ. NSCs were grown as mentioned above. Error Bars are SD. **(G)** Expression of AMPK-α in WT and SR^-/-^ NSCs in control and treated with 0.6 mM L-serine and 0.1 mM D-serine in culture. Actin was a loading control.

### L and D serine rescue defects in *de novo* fatty acid synthesis and differentiation

To determine if treatment with L and D serine can regulate levels of PSPH (final and rate limiting enzyme of the pathway), we treated NSC’s with exogenous L and D-serine (0.6 mM and 0.1 mM respectively). Expression of PSPH was decreased in SR^-/-^ NSCs relative to WT (**Figure 7C**). Treatment with D-serine showed modest increase in PSPH compared to L-serine in SR^-/-^ NSCs and a decrease in WT. These data suggest that availability of both L and D serine regulates expression of PSPH possibly via a feedback mechanism (Fell and Snell, 1988; Snell and Fell, 1990). The magnitude of decrease is greater in WT NSCs while the increased expression in SR^-/-^ NSCs is smaller compared to its own untreated control (**Figure 7C**). Our data on NSCs from SR^-/-^ showed differences in GFAP expression during differentiation (**Suppl Figure 1C**) suggesting that defects in adult neurogenesis in SR^-/-^ mice may result from defects in astrocytic differentiation (Doetsch, 2003). To determine astrocytic differentiation, expression of GFAP was examined under control and rescue conditions. Results show decreased expression of GFAP in differentiating SR^-/-^ NSCs under control condition (**Figure 7A**) compared to WT. Treatment with an equimolar racemic mixture of L and D serine (100 µM each) showed increased GFAP expression in SR^-/-^ and modest increase in WT NSCs (**Figure 7B, Suppl Figure 9C**) suggesting rescue of astrocytic differentiation. Similar rescue in NCAM expression (**Suppl Figure 7A**) was observed in WT and SR^-/-^ NSCs treated with L-serine and D-serine (**Suppl Figures 7B, 7C, 9D**). The magnitude of rescue of NCAM expression appears to be greater in WT and SR^-/-^ NSCs treated with L-serine than D-serine (**Suppl Figure 9D**). These data suggest that both L and D serine can rescue defects in neuroblast formation as evidenced by NCAM expression. To determine if lower levels of malonyl CoA, the precursor for *de novo* lipid biosynthesis in cells can be rescued with similar treatment of NSCs in culture, we estimated levels of malonyl CoA using ELISA. NSC’s from SR ^flox/flox^ and nestin-cre+ grown under control conditions showed approximately 2 fold lower levels in nestin-cre+ compared to age matched SR ^flox/flox^ mice, while treatment with exogenous L and D serine showed 2 fold and higher levels of malonyl CoA in both SR ^flox/flox^ and and nestin-cre+ NSCs which were higher than controls. SR ^flox/flox^ NSCs showed higher levels of malonyl CoA with both L and D serine compared to nestin-cre+. The magnitude of increase appeared to be greater with L-serine treatment than D-serine (**Figure 7D**). These data show both L and D-serine treatment rescue defects in astrocytic, neuronal differentiation and malonyl CoA, with L-serine exhibiting a more potent effect than D-serine.

**Figure 7.**
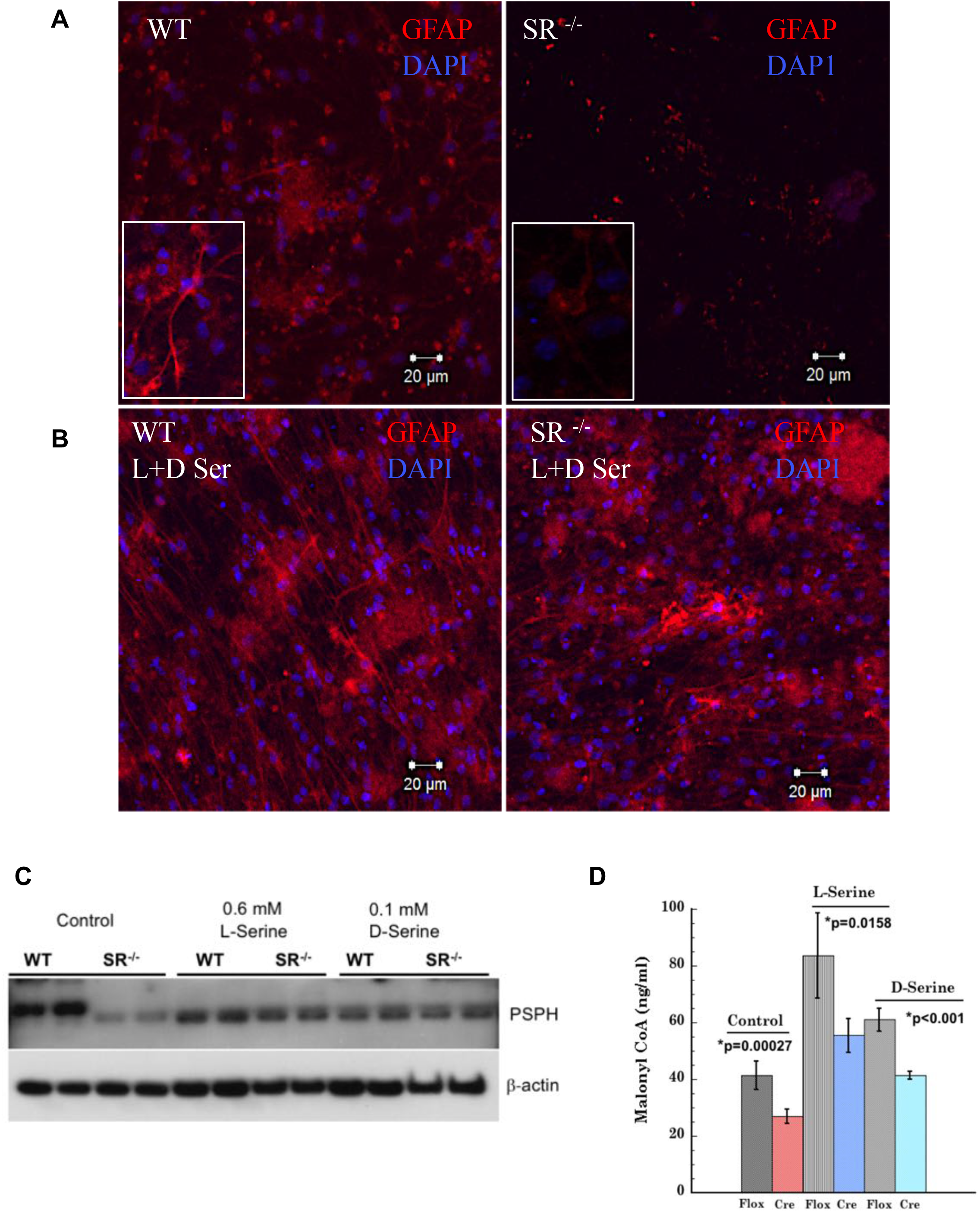
L and D serine rescue defects in *de novo* fatty acid synthesis and differentiation. (**A**) Expression of GFAP (red) in differentiating NSCs from WT and SR^-/-^ mice grown as adherent monolayer culture. (**B**) Expression of GFAP (red) in differentiating NSCs from WT and SR^-/-^ mice grown in presence of equimolar mixture of L-serine and D-serine (100 µM). Scale Bar=20 µ. DAPI is seen in blue. (**C**) Expression of PSPH in WT and SR^-/-^ NSCs treated with L-serine and D-serine. Actin was a loading control. (**D**) Levels of malonyl CoA (ng/ml) in NSCs from SR ^flox/flox^ and nestin-cre+ mice grown in presence of 0.6 mM L-serine and 0.1 mM D-serine. Malonyl CoA was estimated using ELISA. Each group of NSCs was analyzed in triplicate and absorbance at 450 nm corrected from background (540 nm). Amount of malonyl CoA (ng/ml) was quantified from a standard curve. Error Bars are SD.

### Model of SR mediated metabolic control of adult SVZ neurogenesis

We present a simplified summary and model elucidating the role of SR in mediating adult SVZ neurogenesis via lipid metabolism (**Suppl Figure 10**). SR controls the availability of L-serine and D-serine due to its synthesis in astrocytes and neurons respectively (Wolosker et al., 2017) (Ehmsen et al., 2013). Availability of L and D-serine affects the synthesis of different lipids by activation of ACC by an unidentified kinase. Overexpression of Spot14 and Mig12 in nestin-cre+ and SR^-/-^ SVZ regulate the polymerization and catalytic activity of ACC (Knobloch et al., 2013; Park et al., 2013). Activation of ACC by phosphorylation at Ser 79 leads to inhibition of *de novo* fatty acid synthesis by decreasing levels of malonyl CoA and FASN. Reduction in *de novo* fatty acid synthesis affects the formation of newborn neurons in the SVZ (Type A and B cells) and their proliferation, due to loss of their membrane microenvironment (Clemot et al., 2020; Knobloch et al., 2017).

## Discussion

Adult neurogenesis in the SVZ is orchestrated by multiple genes, growth factors and signaling molecules (Kempermann, 2006; Tong and Alvarez-Buylla, 2014). Recent studies have also revealed a metabolic component. Homeostatic energy changes in the brain and in the neurogenic niches appear to influence the proliferative properties of adult neural stem cells. Lipid metabolism plays a major role in adult neurogenesis, both in the hippocampus and SVZ (Clemot et al., 2020; Knobloch et al., 2013; Knobloch and Jessberger, 2017). D-serine functions as a co agonist at the NMDA receptor (Balu et al., 2013; Snyder and Kim, 2000). Aside from this role, its function in the adult brain is unclear. A prior report shows that D-serine mediates adult hippocampal neurogenesis (Sultan et al., 2013) and is involved in proliferation and differentiation of NSCs (Huang et al., 2012). D-aspartate mediates adult neurogenesis in the hippocampus (Kim et al., 2010). D-cysteine, synthesized by SR plays a role in neurodevelopment (Semenza et al., 2021).

We have shown that conditionally deleting SR elicits global changes in lipid classes in the adult SVZ. Our results indicate that SR activates ACC at the primary structural level by phosphorylation at Ser 79 by an unknown kinase (non AMPK dependent) and at the tertiary structural level by affecting polymerization and catalytic activity of ACC due to its interaction with the Spot14 and Mig12 heterocomplex (Knobloch et al., 2013). While L-serine is known to influence cellular proliferation and 1C metabolism in mammals, the role of D-serine in the metabolic process is still unknown.

Lipid rafts are dynamic membrane microdomains enriched in cholesterol and sphingolipids and serve as platforms for relevant signaling cascades involved in stem cell maintenance (Lingwood and Simons, 2010). Our lipidomics data show a bidirectional trend with downregulation of important lipids (ranked by p value) in nestin-cre+ SVZ that affect the generation and proliferation of NSCs. Among the lipid species, a hexosylceramide Hex2Cer40:2;O6 was 3 fold lower while DG O-24:1, a diacylglycerol was 10 fold higher in nestin-cre+ SVZ. SR interacts with PSD95 and stargazin in the activation of AMPA receptor (Ma et al., 2014). Deletion of SR may lead to metabolic consequences due to serine availability and alteration in a variety of signaling pathways. Downregulation of sphingomyelins, ceramides, hexosylceramides and phosphatidylcholines that constitute an integral component of cell membranes was seen in nestin-cre+ SVZ. This downregulation affects neurogenesis and proliferative properties of adult stem cells due to loss of their membrane microenvironment and alterations in lipid raft signaling.

Deletion of SR showed decreased expression of 2 key L-serine synthesizing enzymes, PGDH and PSPH both in the SVZ and NSCs, indicating its compromised synthesis. L-serine is a precursor for the synthesis of phosphatidylserine, sphingosine and ceramide (Esaki et al., 2015). Our lipidomics data reveal that deficiency of L-serine and consequently D-serine in nestin-cre+ SVZ results in 2 fold lower levels of a C22 ceramide species Cer22:0_21:0;O3. Reduction in C16, C18, C20, C22 and C24 acyl chain ceramide species have been reported in mouse embryonic fibroblasts under L-serine deprivation (Esaki et al., 2015). Marked decreases in sphingomyelins, ceramides and hexosylceramides in addition to other classes of lipids were also observed under L-serine deprivation in nestin-cre+ SVZ. Deletion of SR results in significant reduction of D-serine synthesis (Balu et al., 2013; Basu et al., 2009). D-serine levels in both neurons and glia are dependent on PGDH, with lower expression of PGDH leading to decrease in not only L-serine but also D-serine (Ehmsen et al., 2013; Esaki et al., 2015). Altered levels of L and D serine locally may affect cellular metabolism and synthesis of lipid precursors like sphingosine affecting the microenvironment of NSCs. Rescue experiments performed in presence of L-serine, D-serine and a racemic mixture showed metabolic compensation, leading to rescue of proliferation in NSCs, expression of NCAM, Sox2, GFAP and NeuN respectively.

We also observed increased synthesis of malonyl CoA in NSCs grown in presence of L and D serine. Addition of BrdU during differentiation identifies new neurons. SR^-/-^ and nestin-cre+ mice show reduced incorporation of BrdU. Expression of PSA-NCAM, Sox2, and prominin were decreased in nestin-cre+ and SR^-/-^ mice. Endogenous proliferation marker Ki-67 was also decreased in nestin-cre+ SVZ and in SR^-/-^ primary neurons, highlighting their quiescent nature.

While we observed defects in astrocyte and neuronal differentiation, oligodendrocyte differentiation based on expression of Olig2 was enhanced both in SVZ and in NSC’s. Lack of D-serine and or L-serine may favor oligodendrocyte differentiation. Deletion of SR showed protection against cerebral ischemia and excitotoxicity in mice (Mustafa et al., 2010). Our observations highlight a possible role for SR in myelination repair with implications for disorders like multiple sclerosis and in stroke post recovery (Jia et al., 2019).

Hematopoietic stem cells rely on fatty acid oxidation for maintenance and inhibiting this pathway leads to exhaustion of the stem cell pool (Ito et al., 2012). The rationale for investigating lipid metabolism in nestin-cre+ mice arose from the influential role L-serine plays in cell proliferation and the synthesis of sphingolipids. While D-amino acids comprise peptidoglycans in the prokaryotic cell membrane, their function in mammals is unknown. AMPK functions as a master sensor of nutrients and ATP. SR mediates increased phosphorylation of ACC at Ser79 inhibiting synthesis of malonyl CoA, leading to decreased fatty acid synthesis. Interestingly, our data also showed that the activation of ACC was not mediated by AMPK. While nestin-cre+ mice showed increased expression of AMPK and its expression was dependent on L and D serine, it did not undergo activation by phosphorylation at Thr 172 as would be expected. This suggests that the AMPK-ACC activation is not the only axis of control for lipid metabolism. SR may activate additional hubs in an AMPK independent manner. We have yet to identify the kinase responsible for ACC activation. Our data shows that increased expression of Spot14 and Mig12 in nestin-cre+ SVZ may regulate the catalytic activity of ACC due to its effect on ACC polymerization (Knobloch et al., 2013; Park et al., 2013).

Alterations in fatty acid synthesis in the SVZ affects the microenvironment of NSCs and their stemness, resulting in reduced formation of newborn neurons and quiescence. We speculate on few possible connections between serine and lipid metabolism. Glutamate is a known activator of ACC (Brownsey et al., 2006). Due to structural similarities with glutamate, serine may serve as an activator of ACC. PLP is a structural analog of citrate which also regulates ACC. PLP also inhibits both isoforms of ACC (Brownsey et al., 2006). In the absence of SR, excess unbound PLP may inhibit the activity of ACC thereby regulating fatty acid synthesis.

In summary, SR plays a role in lipid mediated control of adult neurogenesis in the SVZ via regulation of ACC activity. Deletion of SR in NSCs affects synthesis of L and D serine with metabolic consequences impacting their proliferative properties. Metabolic control of neurogenesis by administration of L and D-serine may benefit disorders like Alzheimer’s disease and schizophrenia that have an underlying component of adult neurogenesis (Le Douce et al., 2020; Schoenfeld and Cameron, 2015; Yun et al., 2016).

### Experimental Procedures

See supplemental experimental procedures for detailed protocols.

#### Generation of Nestin-Cre+ SR mice

Nestin-Cre+ mice were generated by breeding SR ^flox/flox^ mice (Basu et al., 2009) with nestin cre mice obtained from Jackson labs (number 003771) to generate nestin cre SR flox ^+/-^ or nestin cre (-) SR flox ^+/-^.

#### BrdU (Bromo deoxy Uridine) labeling

Eight weeks old WT and age matched SR^-/-^ mice were kept in a cage and administered 1 mg/ml BrdU (Bromo deoxy Uridine) in the water by means of a water bottle (*ad libitum*) contained within the cage for 14 days.

#### Micro dissection of adult mouse brain SVZ

The mice brain was dissected and removed following euthanasia and kept in a sterile petri dish containing sterile PBS. The petri dish containing the individual brain was placed under a dissecting microscope with light source at low magnification on the ventral surface and the olfactory bulbs were removed by holding the cerebellum. The brain was then rotated on to the dorsal aspect and using a sterile blade a coronal cut was made at the level of the optic chiasm. (Walker and Kempermann, 2014).

#### Adherent monolayer stem cell culture

The neural stem cell adherent monolayer culture system is a valuable tool for determining the proliferative and differentiative potential of adult neural stem cells *in vitro*. Cell culture and differentiation of precursor cells in adherent monolayer cultures was followed as per Walker *et al* protocol (Walker and Kempermann, 2014).

#### Neuroblast Assay

Neuroblasts (Type A cells) were visualized after plating the cells obtained from dissociation of adherent monolayer cultures following accutase treatment. The dissociated monolayer cells were plated in a 15 µ 24 well plate at 2-3 × 10^5^ cells/ml in complete NSC medium supplemented with 20 ng/ml EGF and 10 ng/ml b-FGF. (Azari and Reynolds, 2016; Azari et al., 2012).

#### Neural Stem Cell Immunocytochemistry

Staining of NSC’s for different markers was performed in a 15 µ 24 well plate. The cells were fixed with 4 % PFA in PBS for 20 min at RT and washed once in PBS. Blocking was done in 0.5 % triton X-100 in PBS for 60 min at RT. Following blocking, the cells were incubated with the different primary antibodies in blocking buffer for 60 min at RT.

#### Mouse FASN ELISA

Mouse FASN (fatty acid synthase) expression in mice brain and SVZ region was measured quantitatively using mouse FASN ELISA kit.

#### Mouse Malonyl CoA ELISA

Estimation of mouse malonyl CoA in SVZ lysates was measured quantitatively using mouse malonyl CoA ELISA kit.

#### Quantitative lipidomic analysis

Quantitative lipidomic analysis on SVZ tissue from SR ^flox/flox^ and nestin-cre+ mice were performed at The Metabolomics Innovation Center (TMIC) University of Alberta Canada.

#### Analysis of Gene expression

RNA was isolated from SVZ of WT and SR^-/-^, SR ^flox/flox^ and nestin-cre+ mice using RNA isolation kit. Gene expression studies related to glucose homeostasis and pancreas development were performed using the SYBR green method.

#### Immunohistochemistry

Immunohistochemical experiments were performed on WT, SR^-/-^, SR ^flox/flox^ and nestin-cre+ mice brain coronal and saggital sections. Briefly, the mice were perfused with 4% PFA by cardiac puncture. The brain was removed and incubated at 4°C for 48 h in 4% PFA with gentle shaking. The tissue was paraffin embedded at the Johns Hopkins Oncology Core Services and sectioned at 4 µ thickness.

#### Confocal Microscopy

Fluorescently stained mice brains SVZ sections from age matched WT, SR^-/-^, SR ^flox/flox^ and nestin-cre+ mice were imaged on LSM 780 and LSM 810 confocal microscope (Zeiss) at 10X and 20X magnification at the microscopy core facility at Johns Hopkins. Images were acquired at different high powered fields.

#### Rescue in monolayer adherent cultures

Rescue experiments in adherent monolayer cultures were performed by isolating NSCs from the SVZ of age matched adult WT, SR^-/-^, SR ^flox/flox^ and nestin-cre+ mice. The cells were grown in culture for 7-10 days following which differentiation was induced by gradual removal of growth factors as mentioned in (Walker and Kempermann, 2014).

#### Western blot

Whole brain and SVZ lysates were run on a 1 mm 4-12% Bis-Tris gel with MES SDS running buffer initially at 75 V and then at 120 V. The samples were then transferred to a pre-wet immobilon PVDF transfer membrane and sandwiched between wet filter paper and cassette holder.

#### Time spent sniffing novel odor

Mouse olfactory behavior was tested in age matched WT and SR^-/-^ mice (N=20-25 per group) were collected and brought to the behavior suite and the mice placed in clean cages with new bedding.

#### Blood Glucose Estimation

Blood glucose level was monitored by tail bleeding immediately before and at indicated times after injection using Contour glucometer and Contour blood glucose test strips. Blood glucose measurements were obtained from tail veins at indicated time points post injection.

#### Insulin ELISA

Quantitative estimation of insulin from brain homogenates and SVZ (following microdissection) of age matched WT, SR^-/-^, SR ^flox/flox^ and nestin-cre+ mice were performed using the Ultra Sensitive Mouse Insulin ELISA Kit.

#### Statistics

Statistical tests were computed using KaleidaGraph (Synergy Software, Reading PA). Data are represented as mean ± SD. Unpaired student’s t test was used for pairwise comparison with a fixed control condition. For multiple pairwise comparisons with different control and treatment conditions, one-way ANOVA analysis followed by Tukey’s post hoc test was used. Values with *p* < 0.05 were considered significant.

## Author contributions

**RR** conceived, designed, performed all experiments and analyzed all the data. **RR** wrote the manuscript. **HA** performed neural stem cell differentiation experiments. **SS** reviewed and edited the manuscript. All authors reviewed the manuscript.

## Acknowledgements

The authors acknowledge Ms. Barbara Smith of the Microscopy Core Facility at The Johns Hopkins School of Medicine for confocal imaging. The authors acknowledge Ms. Lauren Albacarys for breeding and performing mice olfactory behavior tests, Dr. Paul Kim for initial technical assistance and Evan R. Semenza of the Snyder laboratory for helpful discussions. The work was funded by grant P50 DA044123 from NIDA to SHS.

