## Supplementary Information for "Serine Racemase mediates Subventricular Zone Neurogenesis via Fatty acid Metabolism"

**1.** Supplemental Figures S1-S11.

**2.** Tables 1-4.

**3.** Experimental Procedures.

**4.** Key Resources Table.

**5.** References.

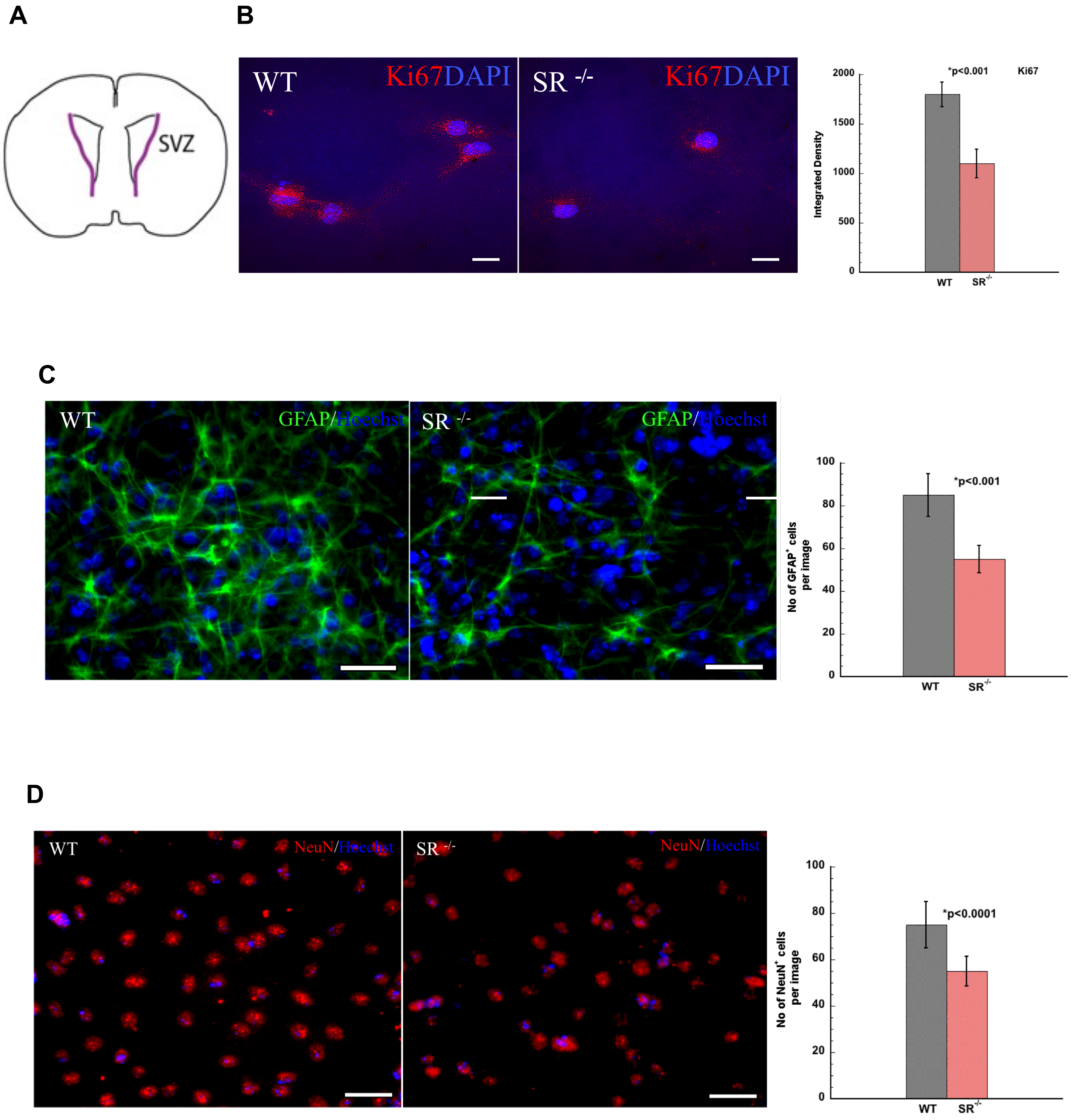

A
SVZ
B
WT
Ki67DAPI
SR -/-
Ki67DAPI
2000
1500
1000
500
0
Integrated Density
*p<0.001
Ki67
WT
SR-/-
C
WT
GFAP/Hoechst
SR -/-
GFAP/Hoechst
100
80
60
40
20
0
No of GFAP+ cells per image
*p<0.001
WT
SR-/-
D
WT
NeuN/Hoechst
SR -/-
NeuN/Hoechst
100
80
60
40
20
0
No of NeuN+ cells per image
*p<0.0001
WT
SR-/-


**Suppl Fig 1.**

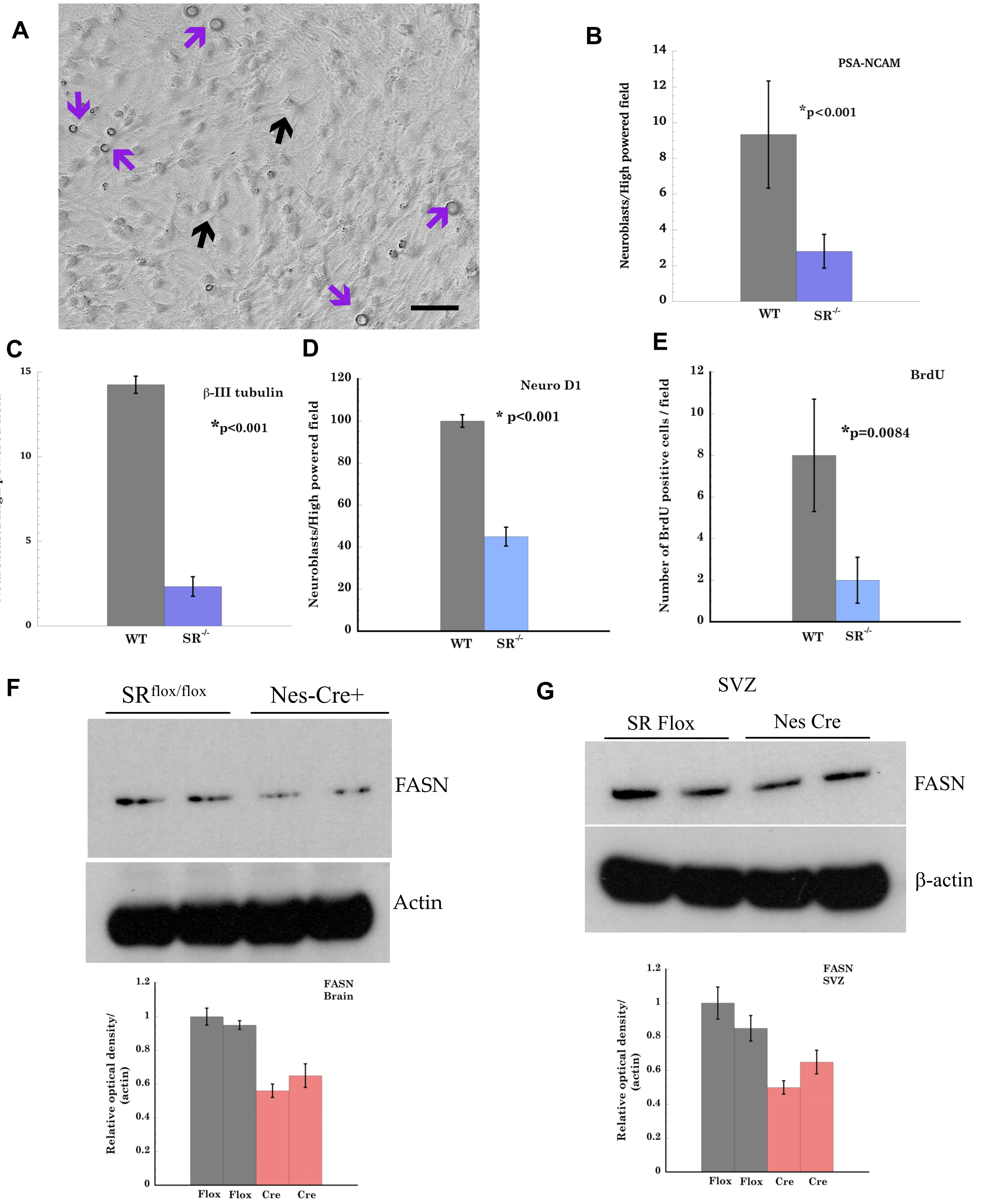
A
B
PSA-NCAM
*p<0.001
Neuroblasts/High powered field
WT
SR-/-
C
β-III tubulin
*p<0.001
Neuroblasts/High powered field
D
Neuro D1
* p<0.001
E
BrdU
*p=0.0084
Number of BrdU positive cells / field
F
SR flox/flox
Nes-Cre+
FASN
Actin
FASN
Brain
Relative optical density/ (actin)
Flox
Cre
G
SVZ
SR Flox
Nes Cre
FASN
β-actin
FASN
SVZ

Suppl Fig 2.

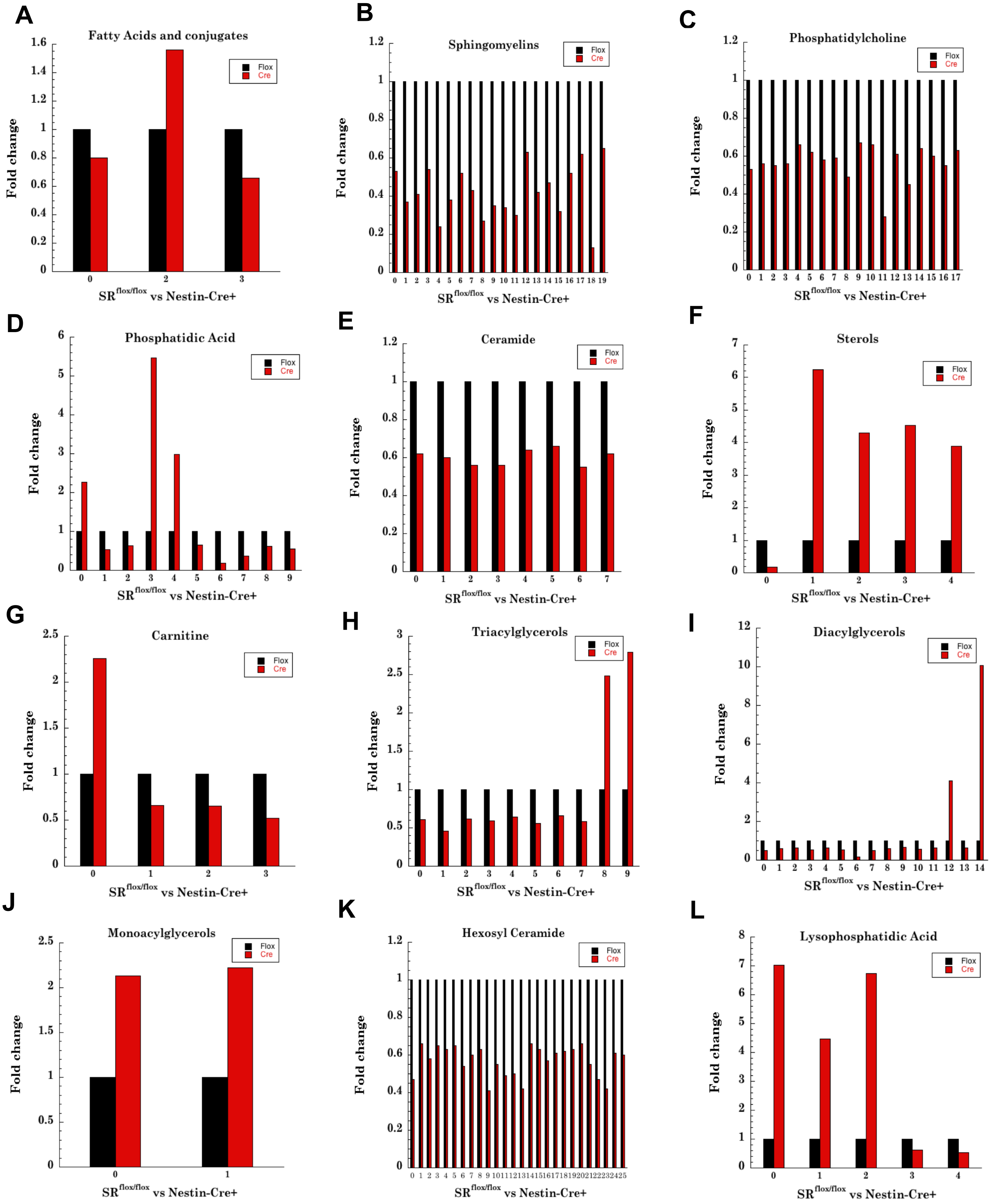
A
Fatty Acids and conjugates
Fold change
SR flox/flox vs Nestin-Cre+
Flox
Cre
B
Sphingomyelins
C
Phosphatidylcholine
D
Phosphatidic Acid
E
Ceramide
F
Sterols
G
Carnitine
H
Triacylglycerols
I
Diacylglycerols
J
Monoacylglycerols
K
Hexosyl Ceramide
L
Lysophosphatidic Acid

Suppl Fig 3.

A

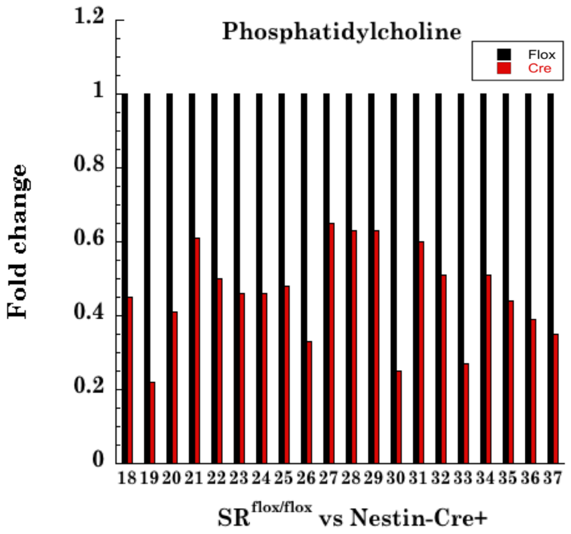


B

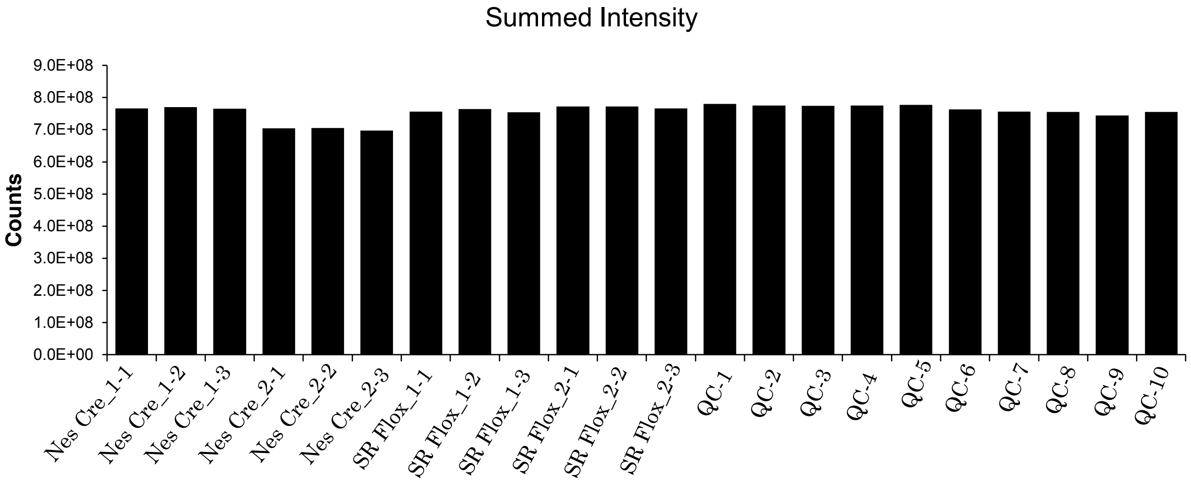


C

| Sample Name | Group | Number of Features |
|---|---|---|
| NesCre_1-1 | Nes Cre | 8257 |
| NesCre_1-2 | Nes Cre | 8231 |
| NesCre_1-3 | Nes Cre | 8274 |
| NesCre_2-1 | Nes Cre | 8181 |
| NesCre_2-2 | Nes Cre | 8162 |
| NesCre_2-3 | Nes Cre | 8156 |
| SR Flox_1-1 | SR Flox | 8202 |
| SR Flox_1-2 | SR Flox | 8224 |
| SR Flox_1-3 | SR Flox | 8184 |
| SR Flox_2-1 | SR Flox | 8186 |
| SR Flox_2-2 | SR Flox | 8204 |
| SR Flox_2-3 | SR Flox | 8196 |
| QC-1 | QC | 8247 |
| QC-2 | QC | 8248 |
| QC-3 | QC | 8247 |
| QC-4 | QC | 8227 |
| QC-5 | QC | 8271 |
| QC-6 | QC | 8192 |
| QC-7 | QC | 8182 |
| QC-8 | QC | 8230 |
| QC-9 | QC | 8227 |
| QC-10 | QC | 8212 |

**Suppl Fig 4.**

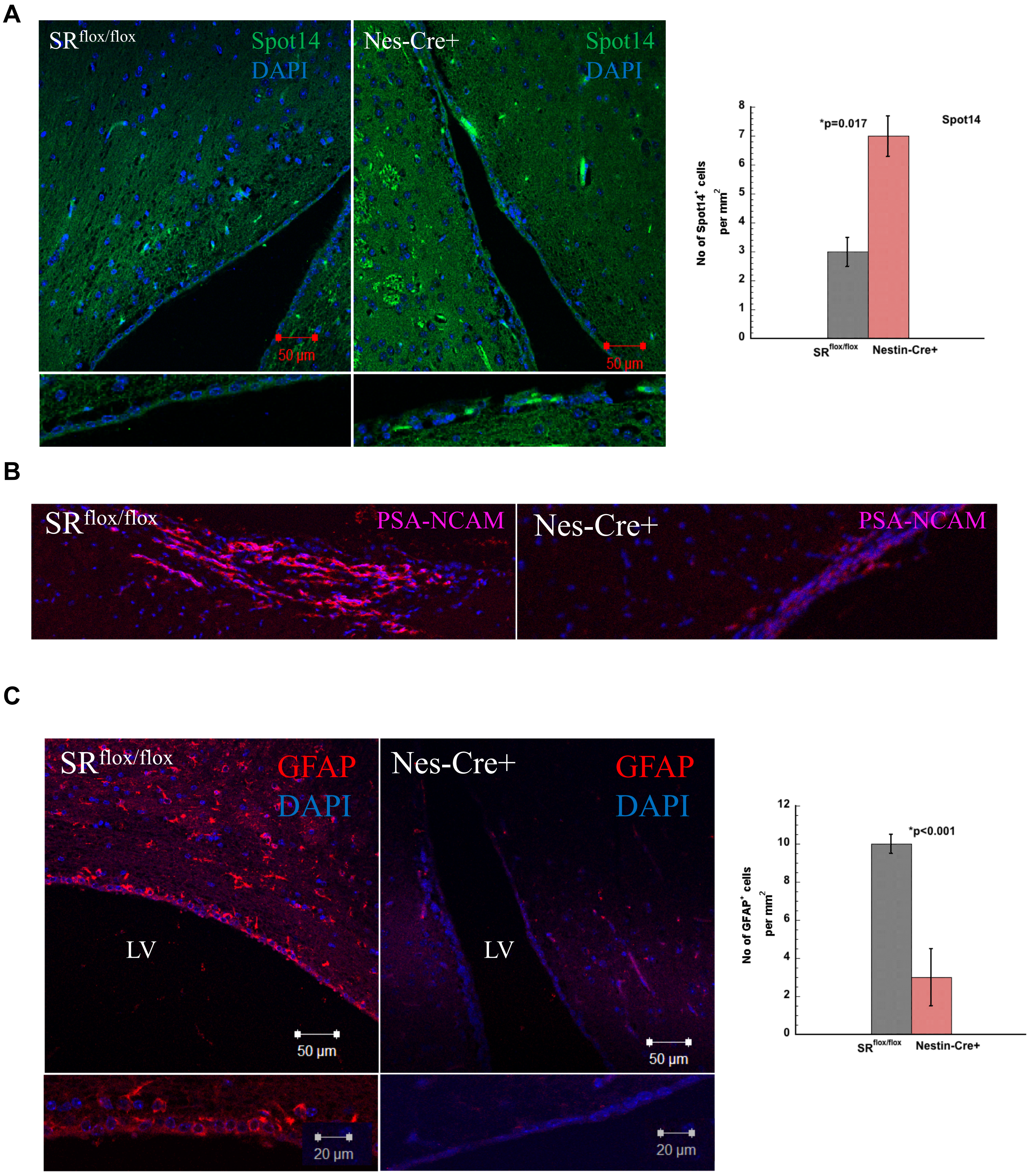
A
SR flox/flox
Spot14
DAPI
Nes-Cre+
Spot14
DAPI
50 µm
50 µm
*p=0.017
Spot14
No of Spot14+ cells per mm2
SR flox/flox
Nestin-Cre+
B
SR flox/flox
PSA-NCAM
Nes-Cre+
PSA-NCAM
C
SR flox/flox
GFAP
DAPI
Nes-Cre+
GFAP
DAPI
LV
LV
50 µm
50 µm
20 µm
20 µm
*p<0.001
No of GFAP+ cells per mm2
SR flox/flox
Nestin-Cre+

**Suppl Fig 5.**

A

B

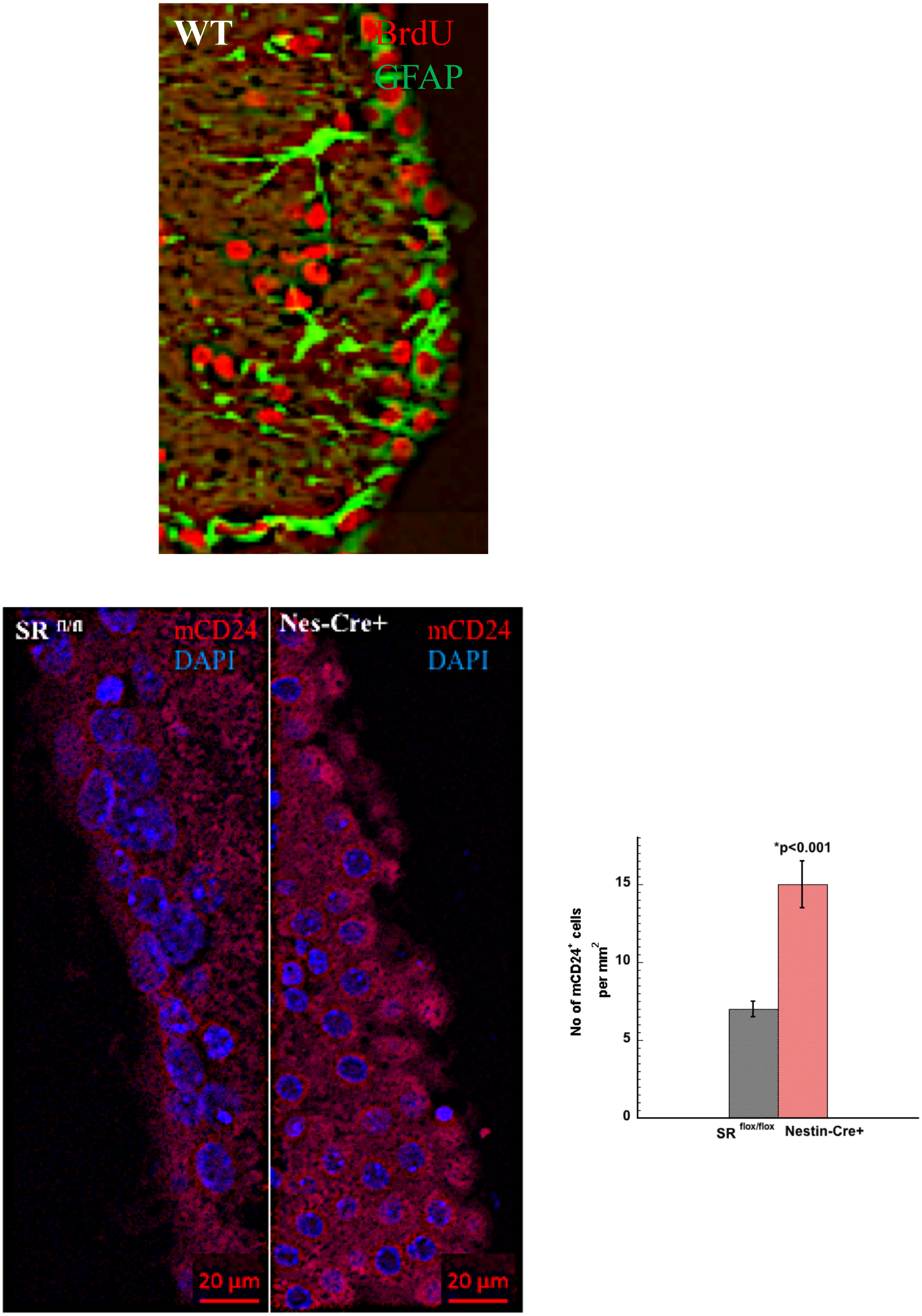


Suppl Fig 6.

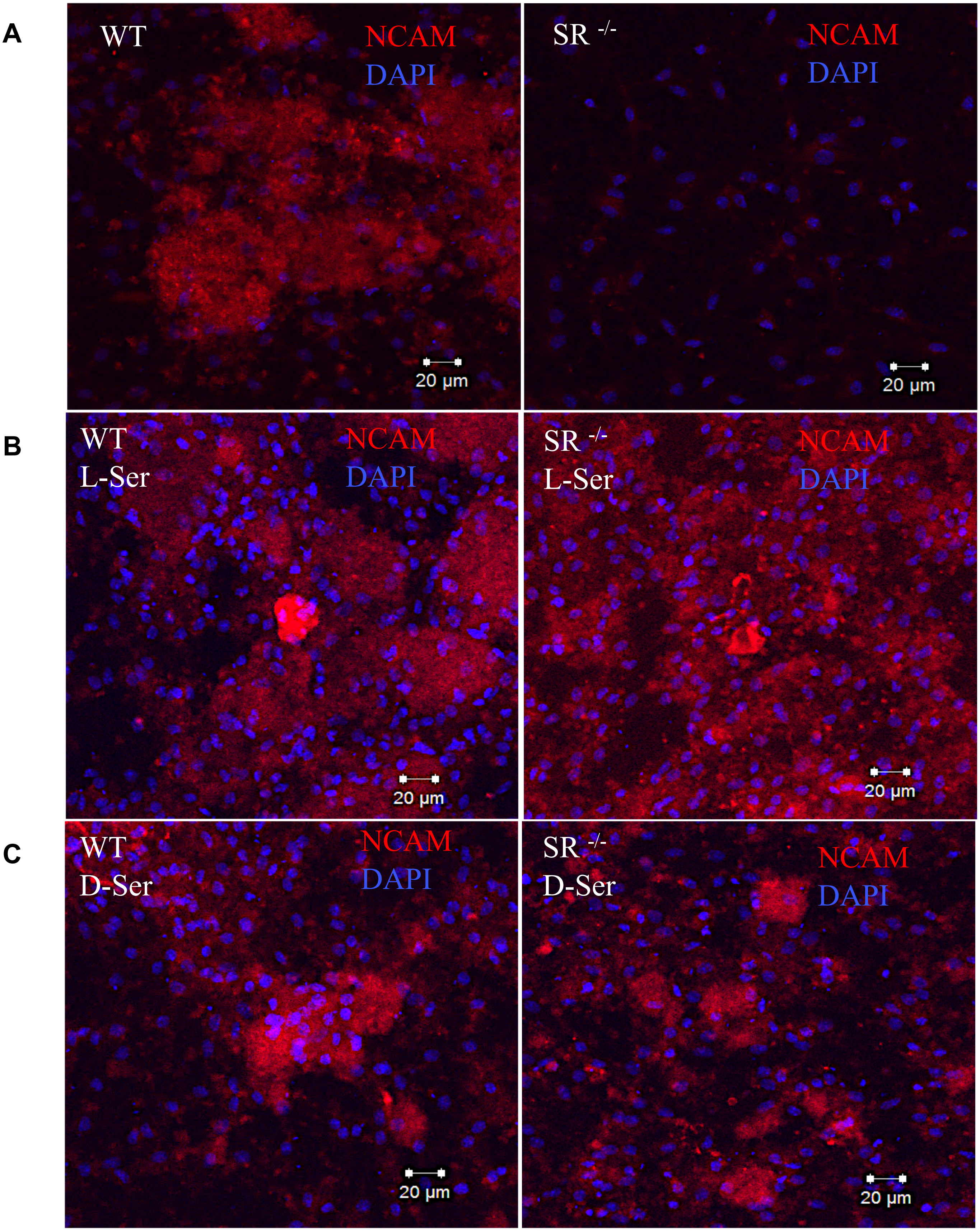
A
WT
NCAM
DAPI
20 µm
SR -/-
NCAM
DAPI
20 µm
B
WT
L-Ser
NCAM
DAPI
20 µm
SR -/-
L-Ser
NCAM
DAPI
20 µm
C
WT
D-Ser
NCAM
DAPI
20 µm
SR -/-
D-Ser
NCAM
DAPI
20 µm

**Suppl Fig 7.**

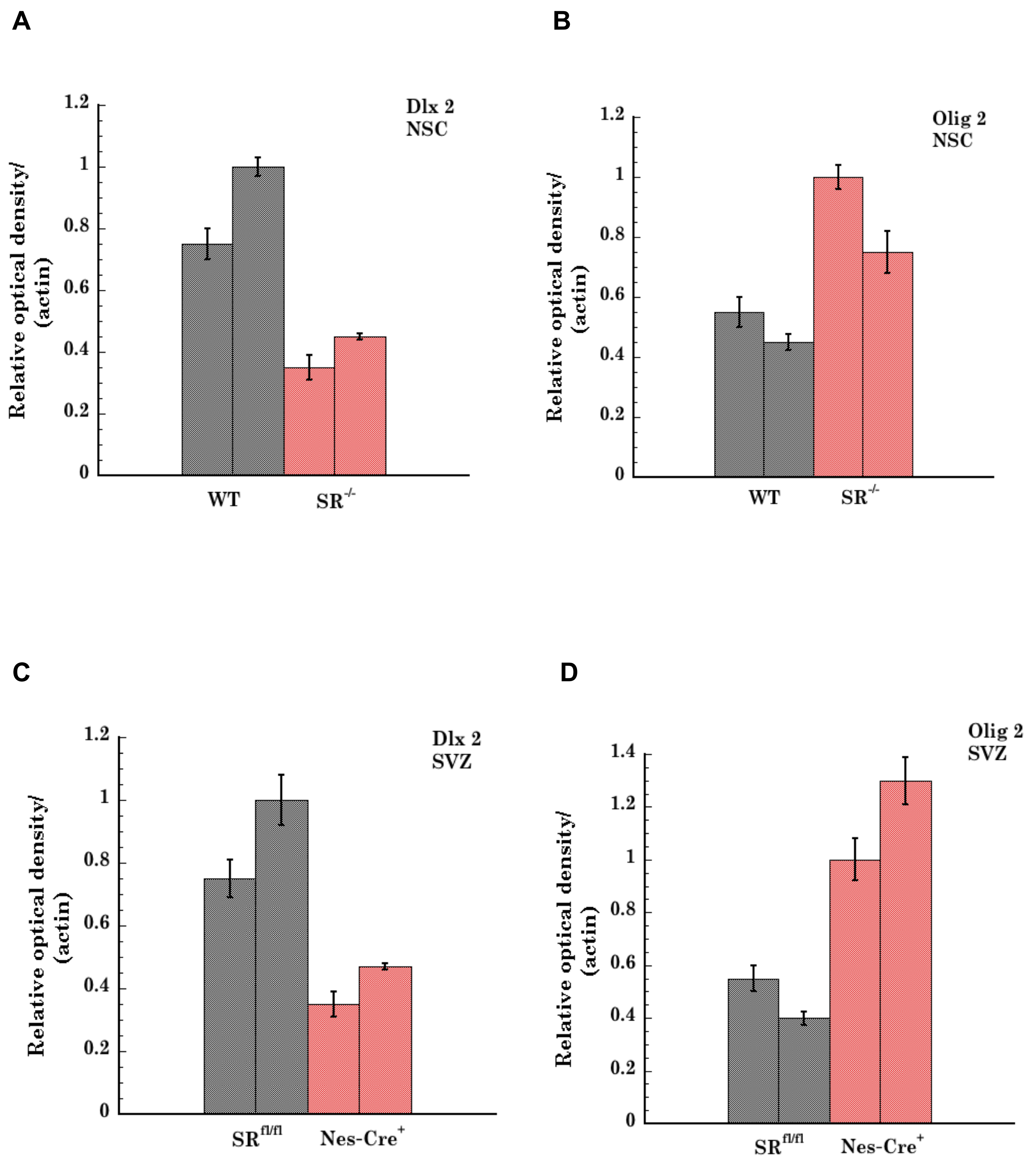
A
Dlx 2
NSC
Relative optical density/
(actin)
1.2
1
0.8
0.6
0.4
0.2
0
WT
SR-/-
B
Olig 2
NSC
Relative optical density/
(actin)
1.2
1
0.8
0.6
0.4
0.2
0
WT
SR-/-
C
Dlx 2
SVZ
Relative optical density/
(actin)
1.2
1
0.8
0.6
0.4
0.2
0
SRfl/fl
Nes-Cre+
D
Olig 2
SVZ
Relative optical density/
(actin)
1.4
1.2
1
0.8
0.6
0.4
0.2
0
SRfl/fl
Nes-Cre+

Suppl Fig 8.

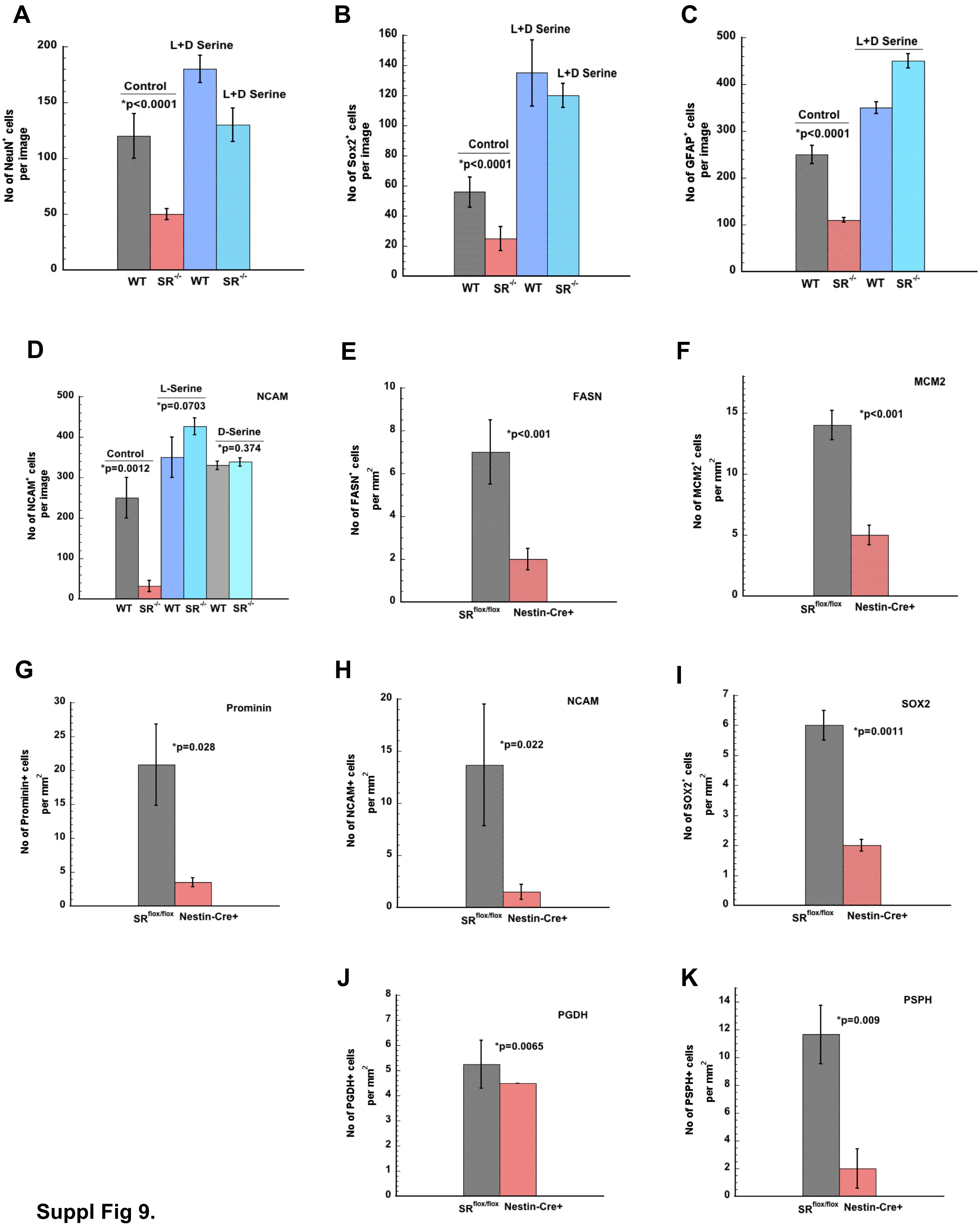
A
No of NeuN+ cells per image
Control
*p<0.0001
L+D Serine
L+D Serine
WT SR-/- WT SR-/-
B
No of Sox2+ cells per image
Control
*p<0.0001
L+D Serine
L+D Serine
WT SR-/- WT SR-/-
C
No of GFAP+ cells per image
Control
*p<0.0001
L+D Serine
WT SR-/- WT SR-/-
D
No of NCAM+ cells per image
NCAM
Control
*p=0.0012
L-Serine
*p=0.0703
D-Serine
*p=0.374
WT SR-/- WT SR-/- WT SR-/-
E
No of FASN+ cells per mm2
FASN
*p<0.001
SRflox/flox Nestin-Cre+
F
No of MCM2+ cells per mm2
MCM2
*p<0.001
SRflox/flox Nestin-Cre+
G
No of Prominin+ cells per mm2
Prominin
*p=0.028
SRflox/flox Nestin-Cre+
H
No of NCAM+ cells per mm2
NCAM
*p=0.022
SRflox/flox Nestin-Cre+
I
No of SOX2+ cells per mm2
SOX2
*p=0.0011
SRflox/flox Nestin-Cre+
J
No of PGDH+ cells per mm2
PGDH
*p=0.0065
SRflox/flox Nestin-Cre+
K
No of PSPH+ cells per mm2
PSPH
*p=0.009
SRflox/flox Nestin-Cre+

Suppl Fig 9.

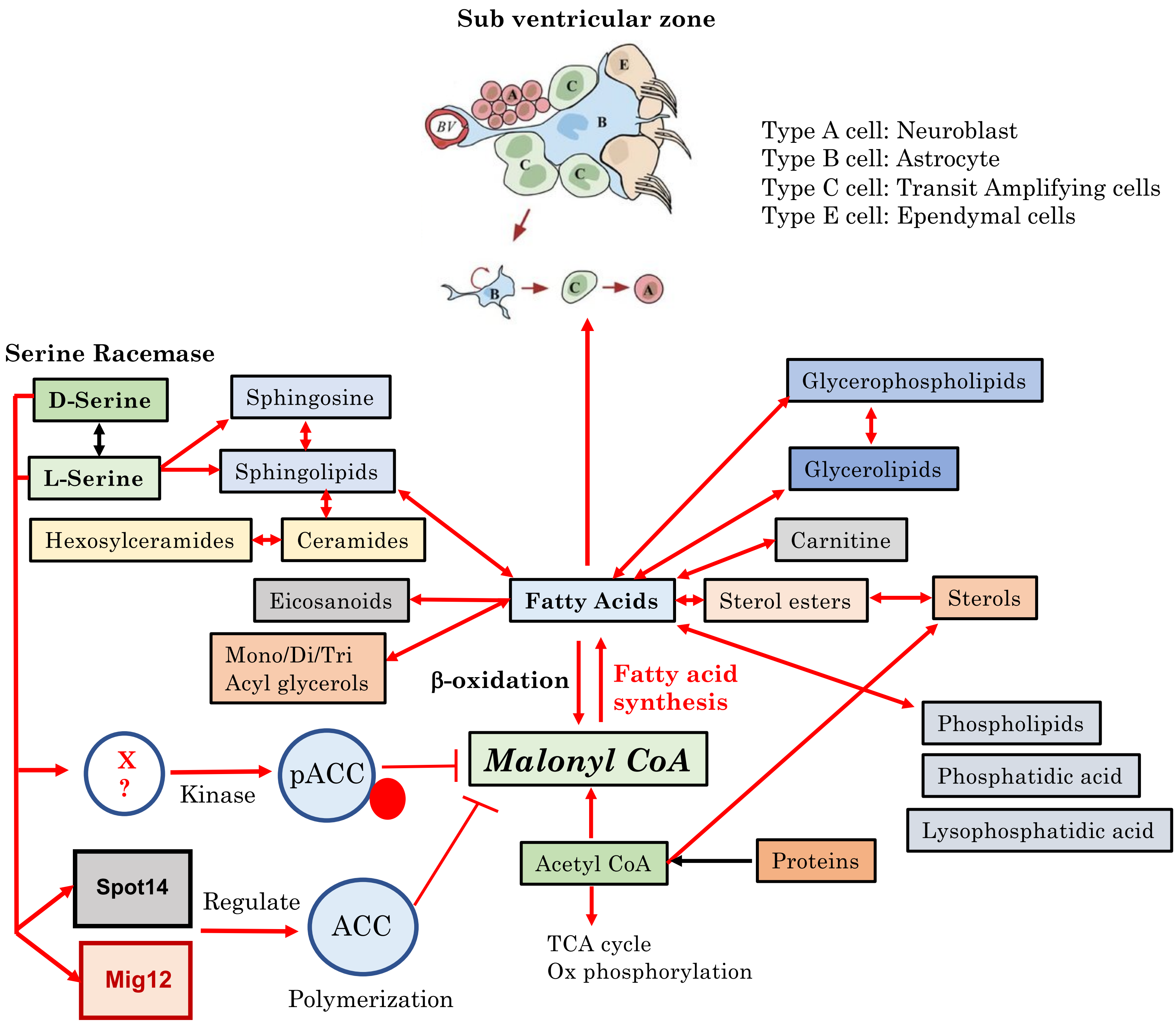

Sub ventricular zone
BV
A
B
C
C
C
E
B
C
A
Type A cell: Neuroblast
Type B cell: Astrocyte
Type C cell: Transit Amplifying cells
Type E cell: Ependymal cells
Serine Racemase
D-Serine
L-Serine
Sphingosine
Sphingolipids
Hexosylceramides
Ceramides
Eicosanoids
Mono/Di/Tri
Acyl glycerols
Fatty Acids
Glycerophospholipids
Glycerolipids
Carnitine
Sterol esters
Sterols
Phospholipids
Phosphatidic acid
Lysophosphatidic acid
β-oxidation
Fatty acid
synthesis
Malonyl CoA
X
?
Kinase
pACC
Spot14
Regulate
ACC
Mig12
Polymerization
Acetyl CoA
Proteins
TCA cycle
Ox phosphorylation


**Suppl Fig 10.**

| Common Name | Abbreviation |
|---|---|
| Acylceramides | Acer |
| Acyl Glucosyl Acylglycerols | AcylGlcADG |
| Bis(monoacylglycero)phosphates (lysobisphosphatidic acids) | BMP |
| Fatty acyl carnitines | Car |
| Cholesteryl esters | CE |
| Ceramides or ceramide phosphates | Cer |
| Cardiolipins | CL |
| Fatty acyl CoEnzyme A's | CoA |
| Diacylglycerol (Diglyceride) | DG |
| Diacylglyceryltrimethylhomoserine | DGTS |
| Ceramide phosphoethanolamines | EPC |
| Fatty acids and conjugates | FA |
| Mono- or digalactosyl mono- or diacylglycerols | GalDG |
| Hexosylceramides | HexCer |
| Ceramide phosphoinositol | IPC |
| Lysophosphatidic acid | LPA |
| Lysophosphatidylcholine | LPC |
| Lysophosphatidylethanolamine | LPE |
| Lysophosphatidylglycerol | LPG |
| Lysophosphatidylinositol | LPI |
| Lysophosphatidylserine | LPS |
| Monoacylglycerol (Monoglyceride) | MG |
| Ceramide phosphoinositols | MIPC |
| N-acyl amines or taurines | NA |
| N-acyl ethanolamines (endocannabinoids) | NAE |
| Others | Other |
| Phosphatidic acid | PA |
| Phosphatidylcholines | PC |
| Phosphatidylethanolamines | PE |
| Phosphatidylglycerols | PG |
| Phosphatidylinositols | PI |
| Phosphatidylinositol-phosphates | PIP |
| Prenols, polyketides, terpenes, flavonoids, and other lipids | PPKT, PR or PK |
| Phosphatidylserines | PS |
| Sulfoquinovosyldiacylglycerols | SDG |
| Sphingomyelins | SM |
| Sphingoid bases or sphingoid base-phosphates | SPB |
| Sterols | ST |
| Sulfoglycosphingolipids (sulfatides) | Sulf |
| Triacylglycerol | TG |

**Suppl Fig 11.**

**Supplemental Figures and Tables legend.**

**Suppl Fig 1. GFAP, NeuN and Ki67 expression in neural stem cells.** (**A**) Schematic of mouse SVZ (**B**) Ki67 staining in WT and $SR^{-/-}$ primary neurons derived from SVZ neural stem cells (**C**) GFAP (**D**) NeuN expression in differentiated WT and $SR^{-/-}$ neural stem cells in culture. DAPI (nuclear stain) is indicated in blue. *Related to Figure 1 and 2.*

**Suppl Fig 2. Quantitative analysis of in vitro adult neurogenesis and expression of mouse FASN.** (**A**) Phase contrast image of neuroblasts (purple arrows) on a bed of astrocytes (black arrows) in the neuroblast assay. Neuroblasts appear 3-4 days after induction of differentiation. Scale Bar=15 μ (**B**) Number of neuroblasts per high powered field in an *in vitro* neuroblast assay in WT and $SR^{-/-}$ mice based on expression of PSA-NCAM (a marker for neuroblast). * indicates $p<0.001$ relative to WT control. Error bars refer to SD. (**C**) Expression of β-III tubulin, a marker for neuroblasts. *$p<0.001$ relative to WT control. Neuroblasts were counted in each of the different high powered fields in each genotype. (**D**) Expression of NeuroD1, a marker for neuroblasts. *$p<0.001$ relative to WT control. (**E**) BrdU (Bromo deoxy Uridine) incorporation in NSC's from WT and $SR^{-/-}$ mice SVZ after incubation for 18 h with 10 μM BrdU. BrdU positive cells were counted in the different high powered fields. *$p=0.0084$ relative to WT control. (**F**) Expression of mouse FASN in whole brain lysates of $SR^{flox/flox}$ and nestin-cre+ mice. (**G**) Expression of mouse FASN in SVZ lysates of $SR^{flox/flox}$ and nestin-cre+ mice. All samples are run in duplicate and pooled from 5-10 mice. Actin was a loading control. Plot below shows densitometry of the blots. *Related to Figures 1 and 3.*

<u>Suppl Fig 3.</u> **Profile of major lipid classes significantly altered in SVZ of SR $^{flox/flox}$ and Nestin-Cre+ mice.** Plots show fold changes in the expression (Y-axis) of lipids in the different classes that are significantly altered in the SVZ of adult SR $^{flox/flox}$ and nestin-cre+ mice following a global lipidomics screen by UPLC-MS/MS. Numbers on the X-axis in the different plots show the specific lipid molecules in the respective classes that are listed in Suppl Table 2. (**A**) Fatty Acid conjugates (**B**) Sphingomyelins (**C**) Phosphatidylcholine (**D**) Phosphatidic Acid (**E**) Ceramides (**F**) Sterols (**G**) Carnitines (**H**) Triacylglycerols (**I**) Diacylglycerols (**J**) Monoacylglycerols (**K**) Hexosyl Ceramides (**L**) Lysophosphatidic Acid. *Related to Figure 3.*

<u>Suppl Fig 4.</u> **List of counts and identified features in LC-MS lipidomics screen** (**A**) Fold change in expression of phosphatidylcholine in the SVZ of SR $^{flox/flox}$ and nestin-cre+ mice from a global lipidomics screen by LC-MS/MS. Numbers on the x-axis indicate individual molecules within the group. The molecules are listed in Suppl Table 4. Y axis indicates fold change in expression relative to SR $^{flox/flox}$ control. (**B**) Summed intensity for all detected features in each LC-MS/MS sample injection in SR $^{flox/flox}$ and nestin-cre+ SVZ after merging positive and negative ionization modes. QC refers to quality control sample. (**C**) Table showing the number of detected features for each sample extraction in SR $^{flox/flox}$ and nestin-cre+ SVZ. QC refers to quality control sample injection which are pooled extracts from both samples. *Related to Figure 3.*

**<u>Suppl Fig 5.</u> Expression of Spot14, PSA-NCAM and GFAP in SVZ.** (**A**) Expression of Spot14 (green) in SR $^{flox/flox}$ and nestin-cre+ mice SVZ along the lateral ventricle. Scale bar=50 μ (top panel). Lower panel shows an enlarged section. (**B**) Expression of PSA-NCAM (pink) along the rostral migratory stream in SR $^{flox/flox}$ and nestin-cre+ mice. Panel shows an enlarged section. (**C**) Expression of GFAP (red) in the lateral ventricle of the SVZ of adult SR $^{flox/flox}$ and nestin-cre+ mice. Lower panel shows an enlarged section of the above figure. Scale bar=50 μ (top panel) and 20 μ (bottom panel). DAPI is indicated in blue. *Related to Figures 1 and 2.*

**<u>Suppl Fig 6.</u> Incorporation of BrdU and expression of mCD24 in SVZ.** (**A**) Representative enlarged section of the lateral ventricle of the SVZ of WT mice showing incorporation of BrdU (red) and GFAP (green). (**B**) Enlarged image of the lateral ventricle of the SVZ in SR $^{flox/flox}$ and nestin-cre+ mice showing expression of mCD24 (red). Scale Bar=20 μ. DAPI is seen in blue. *Related to Figures 1 and 2.*

**<u>Suppl Fig 7.</u> Expression of NCAM in WT and SR$^{-/-}$ NSC's in presence of exogenous L-serine and D-serine** (**A**) Expression of NCAM (red) in NSC's from WT and SR$^{-/-}$ SVZ grown as an adherent monolayer culture for 10 days in untreated condition. (**B**) Expression of NCAM (red) in NSC's from WT and SR$^{-/-}$ grown in presence of 0.6 mM L-serine. (**C**) Expression of NCAM (red) in NSC's from WT and SR$^{-/-}$ grown in presence of 0.1 mM D-serine. Scale Bar=20 μ. DAPI is seen in blue. *Related to Figures 6 and 7.*

**Suppl Fig 8. Densitometric analysis of NSC and SVZ blots.** Densitometric plots of western blots of (**A**) Dlx2 (**B**) Olig2 from NSC lysates of age matched WT and $SR^{-/-}$. Densitometric plots of western blots of (**C**) Dlx2 (**D**) Olig2 from SVZ tissue lysates of age matched $SR^{flox/flox}$ and nestin-cre+ SVZ. Samples were run in duplicate. In each tissue lysate sample, SVZ tissue was pooled from N=5-10 mice. *Related to Figures 1 and 2*.

**Suppl Fig 9. Quantitative analysis of fluorescence in tissue sections and NSC.** Plots show quantitative analysis of (**A**) NeuN (**B**) Sox2 (**C**) GFAP (**D**) NCAM positive staining in rescue experiments performed with SVZ neural stem cells from WT and $SR^{-/-}$ mice. Expression of (**E**) FASN (**F**) MCM2 (**G**) Prominin (**H**) NCAM (**I**) SOX2 (**J**) PGDH (**K**) PSPH in SVZ of age matched $SR^{flox/flox}$ and nestin-cre+ mice. Error Bars refer to SD. * indicates p value between the groups. (student's t-test and ANOVA). *Related to Figures 2, 3, 6, 7 and Suppl fig 7*.

**Suppl Fig 10. Summary schematic of Serine Racemase mediated Acetyl CoA Carboxylase (ACC) control of lipid metabolism**. Simplified schematic of the different lipid classes that are under control of SR in the SVZ of adult mice based on a global lipidomics screen. SR alters lipid metabolism at the primary level by phosphorylation of ACC at Ser 79 and at the tertiary level by altering polymerization of ACC and its loss of catalytic activity by Spot14-Mig12 heterocomplex interaction, leading to decrease in malonyl CoA synthesis and decrease in *de novo* fatty acid synthesis by FASN. SVZ schematic was adapted from (Pircs et al., 2018). pACC (phospho acetyl CoA carboxylase), TCA (tricarboxylic acid). Ox phos (oxidative phosphorylation). *Related to Figure 3, Suppl figs 3 and 4.*

**Suppl Fig 11.** **List of lipid subclass abbreviations.** Table shows the abbreviations of the different lipid subclasses identified in the UPLC-MS MS lipidomics screen. *Related to Figure 3.*

**Suppl Tables Legend.**

**Suppl Table 1.** Table shows the 15 most important lipids identified in the lipidomics screen in the SVZ of age matched adult SR $^{flox/flox}$ and nestin-cre+ mice. Table shows unique identification number, lipid molecule and the class. *Related to Fig. 4F*.

**Suppl Table 2.** Table 2 identities all the lipid molecules from the lipidomics screen in the different lipid classes. Number in each table indicates the lipid molecule identified in the particular class (plotted in suppl fig. 3 on x axis), followed by the lipid ID and fold change in nestin-cre+ mice relative to SR $^{flox/flox}$ (on y-axis). *Related to Fig. 4E*.

**Suppl Table 3.** Table 3 shows the list of primer sequences used for gene expression studies using the SYBR Green method. *Related to Fig. 3B,3I and 5F-G*.

**Suppl Table 4.** Table shows all the lipids identified in the SVZ of SR $^{flox/flox}$ and nestin-cre+ lipidomics heatmap with m/z value, best matched lipid followed by their subclass. *Related to Fig. 4E*.

| Identification # | Lipid | Lipid Class |
|---|---|---|
| P726.5442/11.0 | PE O-22:4_14:0 | Phosphatidylethanolamine |
| N1052.677/16 | Hex2Cer 40:2; O6 | Hexosylceramide |
| N803.5891/13.5 | EPC 40:2; O3 | Ceramide phosphoethanolamine |
| P813.6867/14.1 | SM d24:0_18:2 | Sphingomyelin |
| N723.4963/8.35 | PA 19:2_19:2 | Phosphatidic Acid |
| N913.6994/16.5 | SM 45:3; O3 | Sphingomyelins |
| N956.6584/15.2 | IPC 46:4; O3 | Ceramide phosphoinositol |
| N970.6746/15.7 | IPC 47:4; O3 | Ceramide phosphoinositol |
| P454.3686/2.49 | ST 30:5; O2 | Sterol |
| P458.4204/4.19 | DG O-24:1 | Diacyl glycerol |
| N831.6213/14.7 | EPC 42:2; O3 | Ceramide phosphoethanolamine |
| P815.7027/14.6 | SM d29:0_13:1 | Sphingomyelin |
| N701.5112/10.0 | PA 36:1 | Phosphatidic Acid |
| N984.6901/16.1 | IPC 48:4; O3 | Ceramide phosphoinositol |
| P776.6239/13.4 | HexCer 37:0; O4 | Hexosylceramide |

**Table 1:** Fifteen most important lipids identified from quantitative lipidomics screen in the SVZ of adult SR $^{flox/flox}$ and Nestin-Cre+ mice. Table shows identified lipid followed by lipid class.

**Table 3:** List of primer sequences for SYBR Green method of gene expression.

Spot 14 5'-ATACCAGGAAATGACAGGGCAGGT-3' forward
5'-ACCAAGTCCACAGATGCACTCAGA-3' reverse

Mig12 5'-TTTATCTGTGCTGCTGGAGGAGGT-3' forward
5'-CTGCCCATTCCAGGTGCTCAATTT-3' reverse

Acc1 5'- TTTGCCTTCCGACATCCTGACGTA-3' forward
5'-TCCAGGCTACCATGCCAATCTCAT-3' reverse

Citrate Lyase 5'- TTGGATGCCGCAGCAAAGATGTTC-3' forward
5'- TTGAGGATCTGCACTCGCATGTCT-3' reverse

Hydroxy Acyl CoA (HACoA) 5'-ATGGCGTCCAGTGAGGAGGACGGC-3' forward
5'-TTAATCGTCCTTCTCCGCGATC-3' reverse

Enoyl CoA hydratase (ECH1) 5'- GCTACCGCGATGACAGTTTC-3' forward
5'- TCAGAGATCGAAGGCTGATGTT-3' reverse

Mouse Cpt1a, primers were purchased from BioRad.

Mouse PGDH (MP 201627) and PSPH (MP201727) primer pairs were purchase from Sino Biologicals Inc.

**Experimental Procedures:**

**Reagents:** Purified L-serine and D-serine, were purchased from Sigma Chemical Corp. All salts for buffers and reagents were of research grade and high purity. Milli Q water was used to make buffers and solutions for experiments. HPLC grade solvents and water (Fisher Scientific) was used for estimation of D and L-Serine. Antibodies for rabbit Ki67 (Novus Bio; NB500-170SS), rabbit FASN (Bethyl Labs; A301-324A-T), mouse PSA-NCAM (DSHB Iowa; 5A5-c), mouse Spot14 (sc-137178; Santacruz Biotechnology), rabbit MCM2 (Bethyl Labs; IHC-00009-T), mouse BrdU (Sigma; B8434-25 μl), mouse CD24 (eBioscience; #14-0242-82), rabbit Prominin (Millipore; ZRB1013-25 μl) antibodies were purchased from the vendors mentioned in parenthesis next to the antibody. Antibodies for immunocytochemistry were rabbit NeuN (Cell Signaling Technology; D4G40), βIII-tubulin (RND Systems; #MAB1195-SP), mouse GFAP (Millipore; MAB360), rabbit Olig2 (Novus Bio; 28667SS).

**Generation of Nestin Cre SR mice:** Nestin Cre (conditional deletion of SR) mice were generated by breeding SR flox mice (Basu et al., 2009) with nestin cre mice obtained from Jackson labs (number 003771) to generate nestin cre SR flox $^{+/-}$ or nestin cre (-) SR flox $^{+/-}$. The nestin cre (+) SR flox$^{+/-}$ mice were then bred with SR flox mice to obtain nestin cre SR flox/flox. From the litters obtained and after genotyping (Transnetyx Inc) nestin cre SR flox/flox mice were then bred with SR flox mice to obtain nestin cre SR flox/flox or SR flox/flox mice. The nestin cre SR flox/flox mice were used in experiments along with SR flox/flox mice as controls. All mice were age matched and genotyped before use in experiments (Giusti et al., 2014). Note: Flox and SR Flox refer to SR $^{flox/flox}$ and Nestin Cre to nestin-cre+ (conditional deletion of SR) mice in this study.

**BrdU (Bromo deoxy Uridine) labeling:** Eight weeks old WT and age matched SR$^{-/-}$ mice (and SR $^{flox/flox}$, nestin-cre+ mice) were kept in a cage and administered 1 mg/ml BrdU (Bromo deoxy Uridine) in the water by means of a water bottle (*ad libitum*) contained within the cage for 14 days. The water was changed after 7 days with fresh solution of BrdU. On day 13 the mice were given a single i.p. injection of 200 μl BrdU (5 mg/ml) and the brains harvested after 18 h following euthanasia. The brains were perfused with 4 % PFA in PBS. The brains were sectioned after paraffin embedding. BrdU incorporation in the brain was detected using anti BrdU antibody staining by IHC with antigen retrieval at 60°C for 10 min. The imaging was performed using laser scanning confocal microscopy and a *z* stack obtained. The number of BrdU positive cells were counted in each section and quantified (Doetsch et al., 1999).

**Micro dissection of adult mouse brain SVZ:** The mice brain was dissected and removed following euthanasia and kept in a sterile petri dish containing sterile PBS. The petri dish containing the individual brain was placed under a dissecting microscope with light source at low magnification on the ventral surface and using fine forceps, the olfactory bulbs were removed by holding the cerebellum. The brain was then rotated on to the dorsal aspect and using a sterile blade a coronal cut was made at the level of the optic chiasm. To micro dissect the SVZ, the rostral portion of the brain was positioned to face the cut coronal section upwards. The microscope was focused to a higher magnification. The septum was discarded using fine curved forceps. The SVZ (thin tissue layer adjacent to the ventricle) was dissected by placing the tip of a fine curved forceps in the lateral corner of the ventricle adjacent to the corpus callosum and the other tip approximately 1 mm into the tissue immediately adjacent to the ventricle. The tissue was gently pressed down with the forceps and the triangular piece of tissue removed. The dissected SVZ was placed on a petri dish on ice. A total of 5-10 mice SVZ regions per genotype were pooled in each isolation (Walker and Kempermann, 2014).

**Adherent monolayer stem cell culture:** The neural stem cell adherent monolayer culture system is a valuable tool for determining the proliferative and differentiative potential of adult neural stem cells *in vitro*. The adherent monolayer cultures comprise mostly homogenous population of precursor cells and are useful for following the differentiation process in single cells. The SVZ was carefully dissected from WT, SR$^{-/-}$, SR $^{flox/flox}$ and nestin-cre+ SR mice brains as per the method in (Walker and Kempermann, 2014). The protocol for coating the 96 well and 24 well plate was followed exactly as mentioned. Differentiation of precursor cells in adherent monolayer cultures was followed as per Walker *et al* protocol (Walker and Kempermann, 2014).

**Neuroblast Assay:** Neuroblasts (Type A cells) were visualized after plating the cells obtained from dissociation of adherent monolayer cultures (mentioned above) following accutase treatment. The dissociated monolayer cells were plated in a 15 μ 24 well plate at 2-3 X $10^5$ cells/ml in complete NSC medium supplemented with 20 ng/ml EGF and 10 ng/ml b-FGF. The plated cells were incubated at 37°C incubator for 3-4 days to achieve 75-80 % confluence. Differentiation was induced by gradually replacing media with less growth factors (10 ng/ml b-FGF and 5 ng/ml EGF to no growth factors). The media was subsequently replaced with no growth factors. The differentiated cells (4-5 days post induction) were fixed with 4 % PFA in PBS and immunostained for PSA-NCAM (marker for neuroblasts), β-III tubulin (neuronal marker) and NeuroD1 (neuronal maturation). DAPI was used as a nuclear stain in the mounting media. The cells were visualized under a fluorescence microscope and positively stained cells were counted in each high-powered field and neuroblasts quantified (Azari and Reynolds, 2016; Azari et al., 2012).

**Neural Stem Cell Immunocytochemistry:** Staining of NSC's for expression of different markers was performed in a 15 μ 24 well plate (thin glass bottom; Ibidi; #82406). The cells were washed once with 0.5 ml PBS. The cells were fixed with 4 % PFA in PBS for 20 min at RT. After fixation, the cells were washed once in PBS. The cells were then blocked in 0.5 % triton X-100 in PBS for 60 min at RT. Following the blocking step, the cells were incubated with the different primary antibodies (1:500-1:1000 dilution) in blocking buffer for 60 min at RT. After incubation, the cells washed 3 times in PBS. The cells were incubated with anti-mouse or anti-rabbit Alexa fluor secondary antibodies (1:1000-1:2000 dilution) in blocking buffer for 30 min at RT in the dark. The cells were washed 3 times in PBS. A drop of the mounting medium (ProLong Diamond Antifade Mountant with DAPI; Invitrogen; #P36962) was added to each well and a circular coverslip placed gently in the well. The coverslip was tapped gently to remove any air bubbles. The plate was wrapped in foil and allowed to dry at 4C in the dark. The cells were imaged using a fluorescence microscope.

**Mouse FASN ELISA:** Mouse FASN (fatty acid synthase) expression in mice brain and SVZ region was measured quantitatively using mouse FASN ELISA kit (CUSABIO cat# CSB-EL008435MO). A standard curve was generated from purified mouse FASN as per instructions. Lysates from whole brain and SVZ regions of age matched SR $^{flox/flox}$ and nestin-cre+ mice were homogenized in PBS containing protease inhibitors, sonicated and the harvested supernatant diluted (1:20 v/v) in PBS containing protease inhibitors. Protein concentration was estimated using BCA assay. Equal amounts of protein were added to each well of a 96 well plate in triplicate and the ELISA performed as per manufacturer's instructions. The absorbance at 450 nm was correlated to the amount of FASN (in pg/ml) in the sample based on a FASN standard curve.

**Mouse Malonyl CoA ELISA:** Estimation of mouse malonyl CoA in SVZ lysates was measured quantitatively using mouse malonyl CoA ELISA kit (CUSABIO cat# CSB-E12896m). A standard curve of purified malonyl CoA was generated as per manufacturer's instructions in the concentration range of 0-10 ng/ml. SVZ lysates from age matched SR $^{flox/flox}$ and nestin-cre+ mice were prepared in PBS containing protease inhibitors. The stock lysate was diluted 1:500 in PBS containing protease inhibitors and protein concentration estimated using BCA assay. Equal amounts of protein were added to each well of the 96 well ELISA plate in triplicate. The ELISA was performed as per the instructions in the manufacturer's catalog. Absorbance was read in a microplate reader at 450 nm and background correction of the wells was done at 540 nm. The amount of malonyl CoA (ng/ml) in the SVZ was estimated from the standard curve.

**Quantitative lipidomic analysis:** Quantitative lipidomic analysis on SVZ tissue from SR $^{flox/flox}$ and nestin- cre+ mice (N=5-8 mice pooled per set per genotype; N=2 sets per genotype) and run in triplicate was done at The Metabolomics Innovation Center (TMIC) University of Alberta Canada. The analysis was performed under the four different steps listed below.

**Lipid Extraction.** The extraction was performed strictly following the SOP established based on a modified Folch liquid-liquid extraction protocol. Each sample was extracted in duplicate. The mass of each sample aliquot (5.0 to 6.8 mg) was employed as a normalization factor for the extraction solvents and reagents. Briefly, each aliquot of SVZ brain tissue was vortexed and homogenized with 1.0 µL of internal standard solution / mg of sample and 33.3 µL/mg of methanol, followed by extraction with 65.7 µL/mg of dichloromethane. A clean-up step was performed with 23.6 µL/mg of water. Samples were allowed to equilibrate at room temperature for 10 min and centrifuged at 16,000 g for 10 min at 4°C. An aliquot of the organic layer was evaporated to dryness with a nitrogen blowdown evaporator. The residue was immediately re-suspended in 7.0 µL/mg of NovaMT MixB, vortexed for 1 min, and diluted with 63.0 µL/mg of NovaMT MixA (see below). A pooled mixture of the extracts of both samples was prepared for quality control.

**LC-MS Analysis.** The LC-MS analyses were performed by strictly following the SOP in both positive and negative ionization with extraction duplicates. A total of 12 sample injections (2 samples with extraction duplicates and injection triplicates per genotype) and 10 quality control injections (a pooled mixture of the extracts of both samples) were performed in each ionization polarity. MS/MS spectra were acquired for all samples for identification. Parameters used for data acquisition are described below.

| | |
|---|---|
| **Instrument:** | Thermo Vanquish UHPLC linked to Bruker Impact II QTOF Mass Spectrometer |
| **Column:** | Waters Acquity CSH C18 column, 1.7 µm. |
| **MPA:** | NovaMT MixA |
| **MPB:** | NovaMT MixB |
| **Gradient:** | NovaMT 26-min-gradient |
| **Flow Rate:** | 250 µL/min. |
| **Injection Volume:** | 5.0 µL for positive ionization and 10.0 µL for negative ionization |
| **Column Oven Temperature:** | 42 °C |
| **Mass Range:** | m/z 150-1500 |
| **MS/MS Collision Energies:** | 10-80 eV |

**Data Processing.** LC-MS data from 22 injections were independently processed in positive and negative ionization. Lipid features were extracted and aligned using NovaMT LipidScreener. The data acquired in positive and negative ionization from each sample extraction were combined, i.e. the detected features from all samples were merged into one feature-intensity table. Missing values were substituted by the average intensity of the sample group for features detected in at least 50% of injections within the group (Nestin Cre, SR Flox and QC); or by the global minimum for all sample and QC injections for features detected in less than 50% of injections within the group. Parameters used for data processing are below.

| | |
|---|---|
| Minimum Intensity Cut-off: | 3000 cts for negative ionization; 3000 cts for positive ionization |
| Minimum Peak Length: | 6 spectra |
| Retention Time Tolerance | 5 seconds |
| Mass Tolerance: | 5 mDa |
| Feature Filtering: | Detection for ≥80% of injections in at least one sample group (Nes Cre, SRFlox and QC) |

**Lipid Identification.** A three-tier identification approach based on MS/MS identification and MS match was employed for lipid identification. The parameters used for identification are described below.

| | |
|---|---|
| **Tier 1 (MS/MS identification):** | MS/MS match score ≥ 500; precursor m/z error ≤ 5 mDa |
| **Tier 2 (MS/MS identification):** | MS/MS match score ≥ 100; precursor m/z error ≤ 5 mDa |
| **Tier 3 (MS match):** | Mass match with m/z error ≤ 5 mDa |

After tier 3 identification, a six-tier filtering and scoring approach embedded in NovaMT LipidScreener was employed to restrict the number of matches and select the best identification option to determine the lipid sub-classes for normalization. Data normalization was performed by using a set of 14 deuterated internal standards belonging to different lipid classes. The positively and putatively identified lipids were matched to one of the 14 internal standards according to lipid class similarity and expected retention time range for each class. Intensity ratios, i.e., intensity of each lipid divided by intensity of the matched internal standard, were calculated for normalization. Statistical analysis was performed with MetaboAnalyst 5.0 (https://www.metaboanalyst.ca/). Non-informative features (i.e., internal standards) and features with low repeatability (RSD >20% for QC injections) were filtered out. The dataset was further normalized by auto-scaling and to the median intensity. Finally, the normalized and auto-scaled features were used for statistical analysis.

**Analysis of Gene expression:** RNA was isolated from SVZ of WT and $SR^{-/-}$, $SR^{flox/flox}$ and nestin-cre+ mice using RNA isolation kit (Ambion). Gene expression studies related to glucose homeostasis and pancreas development were performed using the SYBR green method. cDNA synthesis was performed using High Capacity cDNA synthesis kit (Applied Biosystems). 100 ng of synthesized cDNA was used in each well of a 96 well 0.1 ml MicroAmp PCR plate (Applied Biosystems) along with SYBR Green master mix, nuclease free water and gene specific primers (1 $\mu$M final concentration) in a total volume of 10 $\mu$l. 18S rRNA was used as a housekeeping gene and non-template control was added in each plate as a control. Samples were run in triplicate. The PCR was run on an ABI Step One Plus instrument. The raw data obtained was analyzed on Step One data analysis software and mean $C_T$ values obtained. $\Delta C_T$ and $\Delta\Delta C_T$ were calculated and fold change in gene expression obtained from $2^{exp-\Delta\Delta CT}$. The primer sequences spanning exon-exon boundaries for qPCR using SYBR Green are listed in Table 14.

**Immunohistochemistry:** Immunohistochemical experiments were performed on WT, $SR^{-/-}$, SR flox and nestin cre mice brain coronal and saggital sections. Briefly, the mice were euthanized by $CO_2$ narcosis and perfused with 4% paraformaldehyde (PFA) in PBS by cardiac puncture technique. The brain was removed and incubated at 4°C for 48 h in 4% PFA in PBS with gentle shaking. The tissue was paraffin embedded at the Johns Hopkins Oncology Core Services and sectioned at 4 $\mu$ thickness. The sections were de paraffinized using Histoclear (5 min X 2) followed by sequential incubations in absolute alcohol (5 min X 2), 95% alcohol (5 min X 2) and PBS (10 min X 2). The sections were permeabilized in 0.5% Triton X-100 in PBS for 10 min. Antigen retrieval was performed in 10 mM citrate buffer + 0.05% Tween at 60°C for 10 minutes. The sections were cooled on ice for 15 min. The sections were blocked with 1% BSA in PBS for 30 min. The sections were incubated with primary antibody (1:500 dilution in blocking buffer) a 4°C overnight. The slides were washed in PBS (10 min X 2) followed by incubation with secondary antibody (1:1000 dilution in blocking buffer) at RT for 30 min and protected from light. The sections were washed in PBS (10 min X 2). The slides were then cover slipped with mounting medium containing DAPI and edges sealed with transparent nail polish. The slides were allowed to dry at 4°C and observed using confocal microscopy.

**Confocal Microscopy:** Fluorescently stained mice brains SVZ sections from age matched WT, $SR^{-/-}$, $SR^{flox/flox}$ and nestin-cre+ mice were imaged on LSM 780 and LSM 810 confocal microscope (Zeiss) at 10X and 20X magnification at the microscopy core facility at Johns Hopkins. Images were acquired at different high powered fields in each section. All settings and parameters were kept constant during image acquisition. Scale bars were added to each image using Zen black software.

**Rescue in monolayer adherent cultures:** Rescue experiments in adherent monolayer cultures were performed by isolating NSC's from the SVZ of age matched adult WT, $SR^{-/-}$, $SR^{flox/flox}$ and nestin-cre+ mice. Following isolation based on protocol mentioned above, the cells were plated in 500 $\mu$l of growth factor media in coated 24 well plates. After 24 h, the media was replaced completely in each well with fresh media containing either an equimolar mixture of L and D-serine (100 $\mu$M) or individually with L-serine (0.6 mM) and D-serine (0.1 mM). Control cells did not receive any L or D-serine. The media with L and D-serine were replaced every 2-3 days (50 % volume in each well). Both L and D serine were made fresh on the day of addition. The cells were grown in culture for 7-10 days following which differentiation was induced by gradual removal of growth factors as mentioned in (Walker and Kempermann, 2014). All media added to rescue group of cells contained freshly prepared L and D-serine. The cells were harvested 3-4 days after induction of differentiation at which time neuroblasts were present. For immunocytochemistry, the cells were fixed, permeabilized and stained in the 24 well plate. The cells were covered with a coverslip in the well following staining with a drop of antifade mounting medium and stored in the dark at 4°C.

**Western blot**: Whole brain and SVZ lysates were run on a 1 mm 4-12% Bis-Tris gel (Novex Life Technologies; Thermo Fisher) with MES SDS running buffer (Invitrogen; #NP0002-02) initially at 75 V and then at 120 V. The samples were then transferred to a pre-wet immobilon PVDF transfer membrane (Merck Millipore Ltd; Pore Size 0.45 μm) and sandwiched between wet filter paper and cassette holder. The entire apparatus was placed in a wet transfer apparatus (Bio-Rad) and run at 90 V for 90 min on ice. After transfer, the membrane was removed and incubated in blocking buffer containing Tris buffered saline + Tween 20 (TBS-T) containing 5% BSA for 30 minutes at room temperature (RT). The membrane was incubated on a shaker with the respective primary antibody at 1:1000 dilution in blocking buffer overnight at 4°C. After overnight incubation, the membrane was rinsed 3 times with TBS-T and washed in TBS-T at RT for 15 minutes per wash. The membrane was washed a total of 4 times. After washing, the membrane was incubated with HRP conjugated secondary antibody (IgG; mouse or rabbit) (GE Healthcare UK) at 1:5000 dilution in blocking buffer and incubated at RT with shaking for 1 h. After incubation with secondary antibody, the membrane was washed 4 times with TBS-T. The duration of each wash was 15 min. After the last wash, the membrane was incubated with Enhanced Chemiluminescent (ECL) reagent (Thermo Scientific; Super Signal West Pico Plus Peroxide solution and Luminol enhancer solution in a 1:1 (v/v) for 5 min in the dark. The excess ECL reagent was removed using a kimwipe and the membrane was placed in between a plastic sheet and in a cassette holder and developed on an 8” X 10” Ultra Cruz autoradiography film (Santa Cruz Biotechnology) in a developer.

**Time spent sniffing novel odor.** Mouse olfactory behavior was tested in age matched WT and $SR^{-/-}$ mice (N=20-25 per group) were collected and brought to the behavior suite and the mice placed in clean cages with new bedding. The mice were placed in an adjacent room before testing and acclimatized in their new cage for 5 minutes. The new cage had bedding 2-2.5 inches deep. The mice were prepared as mentioned above for this test. Using two disposable pipettes two drops of vanilla extract were placed on one side of the cage and two drops of water were placed on the opposite side of the cage. This eliminates the confounding principle of investigating a novel “object” versus a novel odor. The mice are timed for 2 minutes and recorded for the amount of time it spends sniffing the vanilla extract within the allotted 2 minutes. The data are recorded if the mouse is within 1.5 inches of the placed vanilla extract. The mice are then returned to their original cages. The data were compared between the groups.

**Blood Glucose Estimation.** Blood glucose level was monitored by tail bleeding immediately before and at indicated times after injection using Contour glucometer (Bayer Co, Japan) and Contour blood glucose test strips (Cat#7097C; Ascensia Diabetes care Inc, NJ). Blood glucose measurements were obtained from tail veins at indicated time points post injection. A small drop of blood from the tail was placed on a new glucose strip each time, inserted into the glucometer and value recorded.

**Insulin ELISA:** Quantitative estimation of insulin from brain homogenates and SVZ (following microdissection) of age matched WT, $SR^{-/-}$, $SR^{flox/flox}$ and nestin-cre+ mice was performed using the Ultra Sensitive Mouse Insulin ELISA Kit (Cat# 90080; Crystal Chem). The protocol was followed exactly as mentioned for the low range assay. Standard curve for insulin was developed prior to the actual experiment (low range). Each individual sample was done in quadruplicate. The samples were pooled from N=4-10 mice per genotype and analyzed. The absorbance was read at 450 nm in a 96 well plate reader and also at 630 nm and data plotted after correction at 630 nm.

**Statistics:** Statistical tests were computed using KaleidaGraph (Synergy Software, Reading PA). Data are represented as mean ± SD. Unpaired student's t test was used for pairwise comparison with a fixed control condition. For multiple pairwise comparisons with different control and treatment conditions, one-way ANOVA analysis followed by Tukey's post hoc test was used. Values with $p < 0.05$ were considered significant.

Key Resources Table

| REAGENT or RESOURCE | SOURCE | IDENTIFIER |
|---|---|---|
| Antibodies | | |
| Rabbit anti Ki67 antibody | Novus Bio | NB500-170SS |
| Rabbit anti FASN antibody | Bethyl Labs | A301-324A-T |
| Mouse anti PSA-NCAM antibody | DSHB Iowa | 5A5-c |
| Mouse anti Spot14 antibody | Santacruz Biotechnology | sc-137178 |
| Rabbit anti MCM2 antibody | Bethyl Labs | IHC-00009-T |
| Mouse anti BrdU antibody | Sigma | B8434-25 |
| Mouse anti CD24 antibody | eBioscience | 14-0242-82 |
| Rabbit anti Serine Racemase antibody D5V9Z | Cell Signaling Technology | #29798 |
| Rabbit anti Prominin antibody | Millipore | ZRB1013-25 |
| Rabbit anti NeuN antibody | Cell Signaling Technology | D4G40 |
| Mouse anti β-III tubulin antibody | R&D Systems | MAB1195-SP |
| Mouse anti GFAP antibody | Millipore | MAB360 |
| Rabbit anti Olig2 antibody | Novus Bio | 28667SS |
| Rabbit anti Akt antibody | Cell Signaling Technology | 9272S |
| Rabbit anti phospho-Akt (S473) antibody | Cell Signaling Technology | 4060S |
| Rabbit β-actin antibody | ABClonal | NO:AC026 |
| Goat anti mouse Alexa 547 secondary antibody | Invitrogen | R37121 |
| Goat anti rabbit Alexa 488 secondary antibody | Abcam | ab150077 |
| Bacterial and virus strains | | |

| | | |
|---|---|---|
| N/A | | |
| Biological samples | | |
| N/A | | |
| Chemicals, peptides, and recombinant proteins | | |
| L-serine | Sigma | S4500 |
| D-serine | Sigma | S4250 |
| Ascensia Contour Blood Glucose meter | Bayer Healthcare LLC | 7151B |
| Contour Blood Glucose test strips | Ascensia Diabetes Care | 7097C |
| Critical commercial assays | | |
| Mouse FASN ELISA Kit | CUSABIO | CSB-EL008435MO |
| Mouse Malonyl CoA ELISA Kit | CUSABIO | CSB-E12896m |
| Mouse Insulin ELISA Kit | Crystal Chem | #90080 |
| Deposited data | | |
| N/A | | |
| Experimental models: Cell lines | | |
| N/A | | |
| Experimental models: Organisms/strains | | |
| $SR^{-/-}$ mouse (Serine Racemase knock out mouse) | Basu et al; Molecular Psychiatry 2009 | Joe Coyle, Harvard Medical School |
| WT mouse C57 BL/6 | Jackson Labs | #000664 |
| $SR^{flox/flox}$ mice | | Joe Coyle, Harvard Medical School |

| | | |
|---|---|---|
| Nestin Cre mice (background) | Jackson Labs | 003771 |
| Nestin-Cre+ mice | This paper | |
| Oligonucleotides | | |
| Spot14, Citrate Lyase, Cpt1a, Mig12, Acc primers | Knobloch et al; Nature 2013; Table 3, this paper. | |
| Mouse PSPH qPCR primer pair | Sino Biological | MP201727 |
| Mouse PHGDH qPCR primer pair | Sino Biological | MP201627 |
| HACoA primer | Table 3, this paper | |
| ECH1 primer | Table 3, this paper | |
| Recombinant DNA | | |
| N/A | | |
| Software and algorithms | | |
| Zen Black | Zeiss | https://www.zeiss.com/microscopy/us/products/microscope-software/zen.html |
| Image J | NIH | https://imagej.nih.gov/ij |
| KaleidaGraph v4.1 | Synergy Software | https://www.synergy.com |

| | | |
|---|---|---|
| Slidebook 6 | 3i-Intelligent Imaging Innovations | https://www.intelligent-imaging.com/slidebook |
